# Cell-derived ECM loaded electrospun Polycaprolactone/Chitosan nanofibrous scaffolds for periodontal regeneration

**DOI:** 10.1101/2023.03.30.534964

**Authors:** Mafalda S. Santos, Rachel Cordeiro, Carla S. Moura, Joaquim M. S. Cabral, Frederico Castelo Ferreira, João C. Silva, Marta S. Carvalho

## Abstract

Periodontitis is an inflammatory infection caused by bacterial plaque accumulation that affects the periodontium, a complex structure of different tissues (cementum, periodontal ligament and alveolar bone) that surrounds and supports the teeth. Current treatments lack bioactive signals to induce tissue repair and coordinated regeneration of the periodontium, thus alternative strategies are needed to improve clinical outcomes. Cell-derived extracellular matrix (ECM) has been combined with biomaterials to enhance their biofunctionality for various tissue engineering (TE) applications. In this work, bioactive cell-derived ECM loaded electrospun polycaprolactone/chitosan (PCL/CTS) nanofibrous scaffolds were developed combining polymer solutions with lyophilized decellularized ECM (dECM) derived from human Periodontal Ligament Stem/Stromal Cells (PDLSCs). The work’s aims were to fabricate and characterize cell-derived ECM electrospun PCL/CTS scaffolds in terms of morphology, physico-chemical, thermal and mechanical properties and assess their ability to enhance the osteogenic differentiation of PDLSCs, envisaging periodontal TE applications. PDLSCs were cultured and used for dECM production. PDLSCs-derived dECM was characterized regarding morphology, protein expression, DNA removal efficiency, and glycosaminoglycans and collagen contents. Osteogenic differentiation of PDLSCs was performed on PCL, PCL/CTS and PCL/CTS/ECM electrospun scaffolds for 21 days. The obtained results demonstrate that PCL/CTS/ECM scaffolds promoted cell proliferation compared to PCL and PCL/CTS scaffolds, while maintaining similar physical and mechanical properties of PCL/CTS scaffolds. PCL/CTS/ECM scaffolds enhanced the osteogenic differentiation of PDLSCs, confirmed by increased alkaline phosphatase activity, calcium deposition, and bone-specific marker genes expression. Moreover, PCL/CTS scaffolds showed higher levels of cell mineralization than PCL scaffolds. Overall, this work describes the first use of lyophilized cell-derived ECM loaded electrospun scaffolds for periodontal TE applications and highlights its potential as a promising therapeutic strategy for periodontitis treatment.

## 1. Introduction

Periodontal disease is a chronic inflammatory infection of the periodontium, the structure responsible for ensuring tooth attachment and stability, composed by cementum, periodontal ligament (PDL) and alveolar bone. This infection is caused and sustained by bacteria from dental plaque accumulation [1]. In early stages, periodontal disease manifests as gingivitis, an inflammatory condition of the gingival tissue, which can be reversible with effective oral hygiene. However, if left untreated, gingivitis can progress to periodontitis. Periodontitis, in its advanced form, leads to the degradation and loss of all of the periodontal tissues, characterized by the destruction of collagen fibers, alveolar bone resorption and the formation of soft tissue pockets, causing loose teeth and tooth loss [2]. Current treatments for periodontitis, such as bone grafts and membranes for guided tissue regeneration (GTR), fail to achieve PDL regeneration and integration of soft (PDL) and hard (cementum and alveolar bone) tissues [3]. Current bone grafts mainly serve as a structure for regeneration processes to occur, thus lacking osteoinductivity and osseointegration properties, which are essential for achieving successful bone regeneration [4]. Current GTR membranes have limitations such as low attachment to the adjacent tissues, which can lead to an early exposure of the defect site and allow bacteria infiltration; lack of antibacterial properties and, like bone grafts, poor ability to enhance the regeneration of all the periodontal tissues [5–7]. Alternative strategies to treat periodontitis that lead to a coordinated regeneration of all periodontal tissues are needed to improve clinical outcomes.

Tissue engineering (TE) combines biomaterials, cells, and biologically active chemical/physical factors to develop functional constructs that facilitate tissue regeneration. Periodontal ligament stem/stromal cells (PDLSCs) have been used in periodontal TE strategies. These cells are present in the PDL and are composed of progenitor cells that can self-renew and differentiate into osteoblasts, cementoblasts and fibroblasts, responsible for bone, cementum and PDL formation [8]. PDLSCs have been shown to be able to develop into osteoblasts and cementoblast-like cells *in vitro*, and cementum/PDL-like tissue *in vivo*, and to form collagen fibers connecting to the cementum-like tissue, suggesting their potential to regenerate PDL tissue [9].

Tissue-engineered scaffolds made of synthetic polymers, such as polycaprolactone (PCL), polylactic acid and polyethylene glycol, or of natural polymers, such as chitosan (CTS), gelatin, alginate, collagen and zein, have been studied for periodontal tissue regeneration [10]. Synthetic polymers offer mechanical properties suitable for TE strategies, while natural polymers have high biocompatibility and present bioactive motifs, presenting low cytotoxicity [11]. Electrospinning is a versatile technique to produce fibrous scaffolds for various TE approaches. Electrospun nanofibers can mimic the morphology of extracellular matrix (ECM), therefore facilitating cell attachment, proliferation and differentiation [12]. Electrospinning allows the efficient production of micro/nanofibrous scaffolds that present high porosity, surface area and interconnectivity, which makes these scaffolds highly suitable for the development of biomimetic constructs or periodontal barrier membranes [3]. PCL is a FDA-approved, biodegradable and biocompatible synthetic material that has been extensively used in biomedical applications. PCL can be easily processed, and presents a slow degradation rate (2-3 years) and good mechanical properties suitable for TE [13]. CTS is a natural biodegradable material with antibacterial and osteogenic properties. CTS is obtained through the deacetylation of chitin, which can be extracted from crustaceans’ shells [14]. However, it is highly challenging to produce electrospun CTS fibers alone. Moreover, CTS fibers have poor mechanical properties and rapid degradation [15]. Hence, CTS fibers are usually combined with other polymers, in particular with synthetic polymers like PCL and polyethylene glycol, that provide mechanical support. PCL and CTS have already been used together in strategies for periodontal TE. In different studies, PCL electrospun fibers were coated with drug loaded CTS nanoparticles to serve as vehicles for the administration of antibiotics [16, 17]. Furthermore, blends of both polymers were used to produce PCL-CTS blend fibrous scaffolds. These blend fibers were further incorporated with bioactive glass and hydroxyapatite nanoparticles, resulting in enhanced alkaline phosphatase (ALP) activity [18]. Sundaram and colleagues developed a bilayered construct aiming to simultaneously regenerate alveolar bone and PDL. This construct consisted of PCL electrospun fibers to mimic and regenerate PDL, and a CTS scaffold with calcium sulfate to regenerate the alveolar bone [19].

Recently, decellularized cell-derived ECM (dECM) has been used as a scaffold in several TE strategies [20–23], since it closely mimics the *in vivo*, microenvironment and promotes cell proliferation and differentiation. ECM influences cell proliferation and differentiation through physical, mechanical and chemical cues [24]. Due to its insufficient mechanical properties, particularly when considering hard tissues such as bone, dECM is often combined with biomaterial scaffolds, such as electrospun fibers [25–28] or 3D porous constructs [29, 30], enhancing their biofunctionality for various TE applications. Carvalho and co-workers incorporated dECM in electrospun PCL fibers for bone tissue engineering, which resulted in enhanced cell proliferation and osteogenic differentiation of mesenchymal stem/stromal cells compared to fibers without dECM [25].

In this work, dECM derived from PDLSCs was combined with electrospun PCL/CTS nanofibrous scaffolds. The scaffolds were characterized in terms of their structural, thermal and mechanical properties. Furthermore, osteogenic differentiation of PDLSCs *in vitro*, was evaluated by assessing cell proliferation, calcium production, ECM protein and gene expression. To our knowledge, this is the first study in which dECM derived from PDLSCs was incorporated into PCL/CTS electrospun fibers to develop bioactive and biomimetic ECM loaded nanofibrous scaffolds with good mechanical properties and enhanced bioactivity for improved periodontal TE strategies.

## 2. Materials and Methods

### 2.1. PDLSC Culture

Human PDLSCs used in this work are part of the cell bank available at the Stem Cell Engineering Research Group (SCERG), Institute for Bioengineering and Biosciences (iBB) at Instituto Superior Técnico (IST). Healthy human third molars were extracted for orthodontic reasons from two healthy patients (20 and 28 years old) from the Egas Moniz Dental Clinic, located at Egas Moniz School of Health and Science (Almada, Portugal). The PDL tissue was harvested from the roots of healthy human teeth and PDLSCs were isolated according to established protocols [9]. All human samples were obtained from healthy donors after written informed consent according to the Directive 2004/23/EC of the European Parliament and of the Council of March 31, 2004, on setting standards of quality and safety for the donation, procurement, testing, processing, preservation, storage, and distribution of human tissues and cells (Portuguese Law 22/2007, June 29). Isolated PDLSCs were stored frozen in liquid/vapor nitrogen tanks until further use. PDLSCs were thawed and cultured using low-glucose Dulbecco’s Modified Eagle’s Medium (DMEM, Gibco) supplemented with 10% fetal bovine serum (FBS MSC qualified, Gibco) and 1% antibiotic-antimycotic (A/A, Gibco), and kept at 37°C, 5% CO_2_ in an incubator under humidified atmosphere. Medium renewal was performed every 3-4 days. After reaching confluence, PDLSCs were detached using a 0.05% trypsin solution (Gibco) and counted using the Trypan Blue exclusion method (Gibco).

### 2.2. Decellularized Cell-derived ECM Production

PDLSCs were seeded at a density of 3000 cells/cm^2^ on 6-well plates and culture was maintained with DMEM + 10% FBS + 1% A/A. Cells were expanded for 10 days and medium was renewed every 3-4 days. After 10 days, the medium was discarded and cells were washed with phosphate buffered saline (PBS, Gibco). ECM decellularization was performed using a 0.5% Triton X-100 (Sigma-Aldrich) and 20 mM ammonium hydroxide (NH_4_OH, Honeywell) solution in PBS, based on previously reported methods [20, 31]. PDLSCs were incubated with this solution for 5min at room temperature. After microscopic confirmation of complete cell lysis and presence of ECM on the wells’ surface, dECM was gently washed three times with milliQ water. The dECM was detached using a cell scrapper and collected in falcon tubes. The contents of the falcon tubes were lyophilized to obtain cell-derived dECM powders to be used in the electrospinning procedure.

### 2.3. Characterization of Decellularized Cell-derived ECM

#### 2.3.1. Immunocytochemistry Analysis

The presence and distribution of several ECM proteins in PDLSC culture and PDLSC-dECM were evaluated by immunofluorescence staining of fibronectin, collagen I, laminin, asporin, osteopontin and osteocalcin. Cells and dECM were fixed with 4% paraformaldehyde (PFA, Sigma-Aldrich) for 20 min at room temperature and the immunofluorescent stainings were performed. Samples were washed with 1% Bovine Serum Albumin (BSA, Sigma-Aldrich) for 5 min and then incubated with a blocking solution for 45 min, which was composed of 0.3% Triton-X-100, 1% BSA and 10% FBS in PBS. The primary antibodies, including mouse monoclonal fibronectin (1:400, Abcam), rabbit polyclonal collagen I (1:400, Abcam), rabbit polyclonal laminin (1:400, Abcam), rabbit polyclonal asporin (1:400, Thermo Fisher Scientific), mouse monoclonal osteopontin (1:200, Thermo Fisher Scientific) and mouse monoclonal osteocalcin (1:50, R&D Systems), were added into the samples and incubated overnight at 4°C in a solution of 0.3% Triton X-100, 1% BSA, 10% FBS in PBS. After washing once with 1% BSA, goat anti-mouse IgG-AlexaFluor 546 (1:500, Thermo Fisher Scientific), goat anti-rabbit IgG-AlexaFluor 546 (1:500, Thermo Fisher Scientific) and goat anti-mouse IgG-AlexaFluor 488 (1:500, Thermo Fisher Scientific) secondary antibodies were added into the respective samples and incubated for 1 h at room temperature in the dark. For collagen I, laminin and asporin, the secondary antibody was goat anti-rabbit IgG-AlexaFluor 546; for osteopontin and osteocalcin it was goat anti-mouse IgG-AlexaFluor 546; and for fibronectin it was goat anti-mouse IgG-AlexaFluor 488. Afterwards, samples were washed with PBS and the cells’ nuclei were counterstained with 4’,6-diamidino-2-phenylindole (DAPI, 1.5 μg/mL, Sigma-Aldrich) for 5 min at room temperature in the dark. After washing with PBS, the stainings of the cells and dECM samples were observed and imaged by fluorescence microscopy (Leica DMI3000B).

#### 2.3.2. DNA, sGAG and Collagen Quantification

The amount of DNA, sulphated glycosaminoglycans (sGAGs) and collagens present in samples before and after decellularization was quantified. Cells and lyophilized dECM were stored at −80°C for further use. For each quantification assay, three samples of each condition (N=3) were used. Briefly, the quantification of double-stranded DNA (dsDNA) was measured using a Quant-iT^™^ PicoGreen^™^ dsDNA assay kit (Thermo Fisher Scientific) according to the manufacturer’s instructions. Fluorescence was measured in triplicates on a plate reader (Infinite 200 Pro, Tecan) at an excitation/emission wavelength of 480/520 nm. DNA concentrations of the samples were determined through a standard curve prepared using different concentrations of λ-DNA.

The amount of sGAGs in cells and dECM samples was determined using the 1,9 dimethyl-methylene blue (DMMB) assay according to the manufacturer guidelines. For that, samples were digested at 60°C in a 100 μg/mL Papain (from papaya latex, Sigma-Aldrich) solution (50mM sodium phosphate, 2mM N-acetyl cysteine, 2mM EDTA, all from Sigma-Aldrich, pH 6.5) for 16-18h. Then, samples were combined with the DMMB solution (pH 3.0) for 5 min at room temperature in the dark.Three samples (N=3) were used in the analysis and the absorbance of each sample was measured in triplicates on a plate reader (Infinite 200 Pro, Tecan) at 525 nm. sGAG content was determined through a standard curve using chondroitin-6-sulphate (sodium salt from shark cartilage, Sigma-Aldrich).

The collagen content was quantified using a colorimetric Hydroxyproline assay kit (MAK008, Sigma-Aldrich) following the manufacturer’s instructions. Briefly, cells and dECM samples were hydrolysed with 100 μL of concentrated hydrochloric acid (HCl 12 M, Sigma-Aldrich) for 3h at 120°C. Afterwards, the samples were mixed, centrifuged and the supernatants were transferred to a 96-well plate. The samples were then evaporated to dryness in an oven at 60°C. The assay was then performed through the addition to the samples of ChloramineT/Oxidation Buffer Mixture (5 min incubation at room temperature) followed by a 90 min incubation at 60°C in diluted DMAB reagent. The absorbance of three (N=3) samples for each condition was measured in triplicates on a plate reader (Infinite 200 Pro, Tecan) at 560 nm. The collagen content of the different samples was estimated through a hydroxyproline standard curve.

### 2.4. Fabrication of Cell-derived ECM Electrospun Scaffolds

Polycaprolactone (PCL, Mn = 70,000-90,000 Da, Sigma-Aldrich) was dissolved at 13% w/v in 1,1,1,3,3,3-hexafluoro-2-propanol (HFIP, Tokyo Chemical Industry) under agitation for 2.5 h at room temperature. Medium molecular weight chitosan (CTS, Mn = 190,000-310,000 Da, Sigma-Aldrich) was dissolved at 5% w/v in a Trifluoroacetic acid (TFA, Honeywell)/Dichloromethane (DCM, Honeywell) (70/30 v/v) solvent mixture and stirred for 1.5 h at 50°C, using a magnetic stirrer. PCL and CTS solutions were blended together to obtain a 70/30 v/v PCL-CTS blend solution, followed by agitation overnight. To produce PCL-CTS-ECM electrospun fibers, lyophilized PDLSC-derived ECM was incorporated into the CTS solution (1 mg/mL) and dispersed through agitation for 15 min at 300 rpm using a magnetic stirrer. PCL-CTS and PCL-CTS-ECM scaffolds were crosslinked with glutaraldehyde (GA) vapor (25% v/v, Sigma-Aldrich) for 24 h in a desiccator to ensure their stability. Afterwards, the fibrous scaffolds were fabricated by electrospinning. PCL, PCL-CTS and PCL-CTS-ECM solutions (5 mL) were loaded into a 10 mL syringe placed in a pump and connected to a PTFE tube, which was attached to a 21G stainless steel needle (inner diameter: 0.8 mm). For all solutions, a controlled flow rate of 0.5 mL/h, an applied voltage of 24 kV and a distance of 22 cm between the needle tip and the aluminum foil collector were used. All fibrous scaffolds were electrospun for 2.5 h to ensure scaffold thickness, with temperature and relative humidity varying between 23–24°C and 30–40%, respectively.

### 2.5. Characterization of Electrospun Scaffolds

#### 2.5.1. Scanning Electron Microscopy and Energy Dispersive X-Ray Analysis

Electrospun fibers were characterized morphologically through Scanning Electron Microscopy (SEM) using a Phenom ProX G6 Desktop SEM (Thermo Fisher Scientific). Samples were coated with a gold/palladium layer and imaged at several magnifications, using an accelerating voltage of 10 kV or 15 kV. The average fiber diameters were determined by measuring 100 individual fibers per condition from five different SEM images using ImageJ software (ImageJ 1.51f, National Institutes of Health, USA). The elemental composition of scaffolds was evaluated through Energy Dispersive X-Ray Analysis (EDX) using the Phenom ProX G6 Desktop SEM.

#### 2.5.2. Fourier Transform Infrared Spectroscopy

Attenuated Total Reflectance Fourier Transform Infrared Spectroscopy (ATR-FTIR) analysis was performed using a Bruker AlphaP FTIR spectrometer with Attenuated Total Reflectance platinum–diamond coupling. FTIR spectra were obtained from all of the different fibrous scaffolds and from the individual materials, namely PCL, CTS and dECM, in order to confirm their presence in the electrospun scaffolds. The transmittance spectra of the samples were collected between the spectral region 4000-400 cm^-1^ with a resolution of 4 cm^-1^.

#### 2.5.3. Thermal Properties Analysis

Differential Scanning Calorimetry (DSC) analysis was performed on a Simultaneous Thermal Analyzer, STA 6000 system (Perkin Elmer). Polymers (PCL and CTS) and the different electrospun scaffolds (PCL, PCL-CTS and PCL-CTS-ECM fibers) were weighted in alumina pans (6-8 mg per sample) and heated from 25°C to 200°C at a heating rate of 10°C/min. The thermal degradation of the polymers and the fibers was evaluated using Thermogravimetric Analysis (TGA) mode on the STA 6000 system. The samples were heated from 50°C to 600°C at a heating rate of 10°C/min. DSC and TGA were performed in triplicates under a nitrogen atmosphere with a flow rate of 20 mL/min.

#### 2.5.4. Contact Angle measurements

The contact angles of PCL, PCL-CTS and PCL-CTS-ECM fibers were measured using a DSA25 Drop Shape Analyzer (Krüss) in the sessile drop method. Droplets of distilled water were placed on the scaffolds’ surface and the contact angles were measured. For each condition, the contact angles were measured in three (N=3) individual fiber samples and the results were analyzed using the software Drop Shape Analysis 4 version 2.1.

#### 2.5.5. Mechanical Tensile Testing

The mechanical properties of electrospun scaffolds were assessed through uniaxial tensile testing using a mechanical tester (Univert Model UV-200-01, CellScale Biomaterials Testing), with a 10 N load cell and a displacement rate of 3 mm/min. For each condition, ten different test specimens (N=10) were cut as rectangular strips (with a length of 30 mm, width of 10 mm and thickness of 0.1mm) and tested. The experimental data was collected and processed using the UniVert software. The Young’s modulus of each specimen was obtained from the slope of the initial linear strain region (0-15%) of the stress-strain curve. The ultimate tensile strength (UTS) and ultimate elongation were also obtained from the stress-strain curves.

### 2.6. In Vitro Cell Culture on Electrospun Scaffolds and Assessment of their Biological Effects

#### 2.6.1. Scaffold preparation and cell seeding

Prior to cell culture, PCL, PCL-CTS and PCL-CTS-ECM electrospun scaffolds were sterilized with UV light for 30 min and washed three times (1 h each wash) with PBS + 1% A/A solution. The scaffolds were placed in ultra-low cell attachment 24-well plates, washed again with PBS + 1% A/A, and then incubated in culture medium for 1 h at 37°C. PDLSCs were seeded on the different electrospun scaffolds at a density of 50,000 cells per scaffold and incubated for 2 h at 37°C and 5% CO_2_ to promote initial cell attachment. Osteogenic medium, composed of DMEM supplemented with 10% FBS, 1% A/A, 10 mM β-glycerophosphate (Sigma-Aldrich), 50 μg/mL ascorbic acid (Sigma-Aldrich) and 10 nM dexamethasone (Sigma-Aldrich) was added to all scaffolds. PDLSCs were cultured on the scaffolds for 21 days and medium was renewed every 3-4 days.

#### 2.6.2. PDLSC Viability and Proliferation Assay

The metabolic activity of PDLSC in the different electrospun scaffolds was evaluated using AlamarBlue^®^ Cell Viability Assay (Thermo Fisher Scientific) on days 1, 7, 14 and 21. A 10% (v/v) AlamarBlue^®^ solution prepared in culture medium was added to the scaffolds and incubated at 37°C in 5% CO_2_ chamber for 3 h. The fluorescence was then measured on a plate reader (Infinite 200 Pro, Tecan) at an excitation/emission wavelength of 560/590 nm and compared to a calibration curve to assess the equivalent number of cells in each scaffold. Six independent scaffolds (N=6) were used for each condition and fluorescence was measured in triplicates. Scaffolds without cells were used as blank controls.

#### 2.6.3. Cell Morphology

To assess the morphology of PDLSCs present on the electrospun scaffolds, a DAPI-Phalloidin staining was performed. Cells were washed with PBS, fixed with 4% PFA for 20 min and permeabilized with a 0.1% Triton X-100 solution for 10 min. Afterwards, samples were incubated with Phalloidin-TRITC (1:100 in PBS, Thermo Fisher Scientific) for 45 min at room temperature in the dark. Then, cells were washed with PBS, counterstained with 1.5 μg/mL DAPI solution for 5 min and washed with PBS. The fluorescent stainings were imaged by fluorescence microscopy (Leica DMI3000B).

The cell morphology of the PDLSCs cultured on the electrospun scaffolds was also analyzed using SEM (Phenom ProX G6 Desktop SEM). After 21 days of osteogenic differentiation, scaffolds were washed once with PBS, fixed with 4% PFA for 20 min and dehydrated using ethanol gradient solutions (20%, 40%, 60%, 80%, 90% and 100% (v/v), 30 min incubation in each solution). Then, the samples were incubated for 1 h in hexamethyldisilazane (HDMS) and left to air dry inside a fume hood. Scaffolds were coated with a gold/palladium layer and then analyzed using SEM as previously specified in section 2.5.1.

#### 2.6.4. Energy Dispersive X-Ray Analysis

Aiming to evaluate the cell mineralization after 21 days of PDLSC osteogenic differentiation on the different electrospun scaffolds, EDX analysis was performed with an accelerating voltage of 10 kV, according to section 2.5.1.

#### 2.6.5. ALP/Von Kossa Stainings

After 21 days under osteogenic differentiation conditions, the scaffolds were washed once with PBS, fixed with 4% PFA for 20 min and stained with ALP and Von Kossa stainings to identify ALP activity and mineral deposits, respectively. First, the scaffolds were incubated with a Fast Violet solution (Sigma-Aldrich) and Naphthol AS-MX Phosphate Alkaline solution (Sigma-Aldrich) in a final concentration of 4% for 45 min at room temperature, in the dark. For the Von Kossa staining, the same scaffolds were incubated with a 2.5% silver nitrate solution (Sigma-Aldrich) for 30 min at room temperature in the dark. Finally, the samples were washed twice with milliQ water and imaged under the microscope.

#### 2.6.6. ALP Activity Quantification Assay

ALP activity was quantified using QuantiChrom ALP Assay Kit (BioAssays Systems) according to the manufacturer’s protocol after 21 days of PDLSC osteogenic differentiation on the different types of nanofibrous scaffolds. Briefly, the samples were washed with PBS and were incubated with a cell lysis solution (0.1% Triton X-100 in PBS) overnight at room temperature under orbital agitation. Then, a 10 mM p-nitrophenyl phosphate solution, provided with the ALP kit, was added to the samples. The absorbance was measured on a plate reader (Infinite 200 Pro, Tecan) at 405 nm and normalized to the total number of cells in each scaffold. For each condition, three different scaffolds (N=3) were used for each condition and the absorbance values were collected in triplicate.

#### 2.6.7. Alizarin Red Staining and Quantification

After 21 days under osteogenic differentiation conditions, calcium deposition was assessed and quantified by Alizarin Red staining. For each condition, three scaffolds (N=3) were washed once with PBS, fixed with 4% PFA for 20 min and incubated with a 2% Alizarin Red solution (Sigma-Aldrich) for 1 h at room temperature, in the dark. Then, the scaffolds were washed several times with milliQ water and imaged under the microscope. Afterwards, the Alizarin Red was quantified by dissolving it with a 10% (w/v) cetylpyridinium chloride solution under agitation for 1h. The absorbance was measured in triplicates on a plate reader (Infinite 200 Pro, Tecan) at 550 nm and compared to a standard curve to determine Alizarin Red concentration.

#### 2.6.8. Biomineralization (OsteoImage) Assay

PDLSCs mineralization on the scaffolds after 21 days of osteogenic differentiation was assessed using OsteoImage Mineralization Assay (Lonza). This assay’s staining reagent binds specifically to the hydroxyapatite portion of bone-like nodules deposited by cells. For each experimental group, three scaffolds (N=3) were washed once with PBS, fixed with 4% PFA for 20 min and washed once with 1X Wash Buffer. Then, the samples were incubated with a diluted staining reagent for 30 min at room temperature, in the dark. Scaffolds were then washed with 1X Wash Buffer and the fluorescent staining was observed by a fluorescence microscopy (Leica DMI3000B). Furthermore, the fluorescence intensity values were measured on a plate reader (Infinite 200 Pro, Tecan) at an excitation/emission wavelength of 492/520 nm.

#### 2.6.9. RNA Extraction and qRT-PCR Analysis

Total RNA was extracted using the RNeasy Mini Kit (QIAGEN). Briefly, the scaffolds were first incubated in lysis buffer for 1 h with agitation. Then, total RNA was isolated following the manufacturer’s protocol and quantified using a NanoVue Plus spectrophotometer (GE Healthcare). cDNA was synthesized from the purified RNA using High-Capacity cDNA Reverse Transcription kit (Applied Biosystems). Reaction mixtures were incubated in a T100^™^thermal cycler (Bio-Rad) for 5 min at 25°C, 20 min at 46°C and 1 min at 95°C and then were maintained at 4°C. The quantitative reverse transcription-polymerase chain reaction (qRT-PCR) was performed using NZYSpeedy qPCR Green Master Mix (2x), ROX plus (NZYTech) and the equipment StepOne Plus real-time PCR system (Applied Biosystems) following the manufacturer’s guidelines. All samples were analyzed in triplicates (N=3) and using the 2-ΔΔCT method. Target gene expression was primarily normalized to glyceraldehyde 3-phosphate dehydrogenase (GAPDH) gene expression and then determined as a fold-change relative to the baseline target gene’s expression in undifferentiated PDLSCs at day 0, prior to scaffold seeding (Control). The target genes analyzed were Runt-related transcription factor 2 (RUNX2), Osterix (OSX), osteocalcin (OC), alkaline phosphatase (ALP), collagen I (COL I) and cementum protein 1 (CMP1). Primer sequences used in the qRT-PCR analysis are presented in Table 1.

**Table 1:**
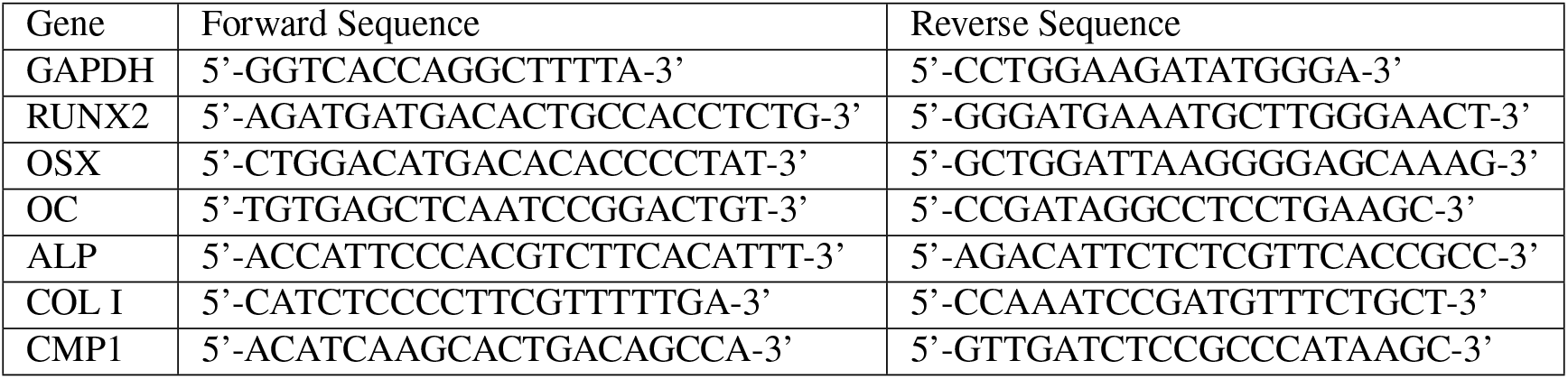
Sequences of primers used for qRT-PCR analysis.

#### 2.6.10. Immunocytochemistry Analysis

The presence of collagen I (COL I), asporin (ASP), osteopontin (OPN), osteocalcin (OC), periostin (POSTN) and cementum protein 1 (CMP) was analyzed in PDLSCs cultured on electrospun scaffolds for 21 days under osteogenic differentiation conditions. The scaffolds were washed once with PBS and fixed with 4% PFA for 20 min at room temperature. Cells were washed with a 1% BSA solution for 5 min and then samples were blocked with a solution of 0.3% Triton X-100, 1%BSA, and 10% FBS (in PBS) at room temperature for 45 min. The primary antibodies, including rabbit polyclonal collagen I (1:200, Abcam), rabbit polyclonal asporin (1:100, Thermo Fisher Scientific), mouse monoclonal osteopontin (1:100, Thermo Fisher Scientific), mouse monoclonal osteocalcin (1:50, R&D Systems), rabbit polyclonal periostin (1:100, Abcam) and rabbit polyclonal cementum protein 1 (1:100, Thermo Fisher Scientific) prepared in a solution of 0.3% Triton X-100, 1% BSA, 10% FBS (in PBS) were added into the samples and incubated overnight at 4°C. After overnight incubation, the samples were washed once with 1% BSA and the secondary antibodies were added into the respective samples and incubated for 1 h at room temperature, in the dark (goat anti-rabbit IgG Alexa Fluor 546 [1:200, Thermo Fisher Scientific] and goat anti-mouse IgG Alexa Fluor 546 [1:200, Thermo Fisher Scientific]). For collagen I, asporin, periostin and cementum protein 1, the secondary antibody was goat anti-rabbit IgG-AlexaFluor 546; and for osteopontin and osteocalcin it was goat anti-mouse IgG-AlexaFluor 546. Finally, scaffolds were washed with PBS and the cells’ nuclei were counterstained with 1.5 μg/mL DAPI solution (Sigma-Aldrich) for 5 min at room temperature in the dark. After washing with PBS, the immunofluorescent stainings were observed and imaged by fluorescence microscopy (Leica DMI3000B).

### 2.7. Statistical Analysis

The statistical analysis of the data was performed in GraphPad Prism 9 software using one-way ANOVA, followed by Tukey post-hoc test. Data was considered to be statistically significant when the p-values were less than 0.05 (95% confidence intervals, *p < 0.05).

## 3. Results

### 3.1. Characterization of decellularized ECM derived from PDLSCs

Confluent cultures of PDLSCs were decellularized according to previously published procedures [20, 25, 31]. Before decellularization, PDLSCs stained positive for fibronectin, collagen I, laminin, asporin, osteopontin and osteocalcin (Figure 1A). After decellularization, dECM derived from PDLSCs retained the expression of these proteins (Figure 1A). Furthermore, DAPI staining demonstrated only a residual amount of cellular nuclei, indicating that most of the cellular nuclei were removed. dECM obtained after decellularization presented a fibrillar structure staining positive for collagen I, laminin, and fibronectin. Additionally, PDLSCs and dECM were imaged under bright field microscopy and a DAPI-Phalloidin staining was performed, as described in section 2.6.3 (Supplementary Figure A.1). Bright field images show that PDLSCs were confluent before decellularization (A.1A) and that the obtained dECM presented a fibrillar structure (A.1B). Before decellularization, PDLSCs possessed well-defined cell nuclei and cytoskeleton (A.1C), whilst after decellularization no cell nuclei or actin filaments were visible (A.1D). To confirm an efficient decellularization, DNA quantification was evaluated after the decellularization treatment. Results showed that the DNA content of the dECM was significantly diminished after the decellularization process as expected (Figure 1B). After decellularization, dECM retained to some extent the sGAGs and collagen content present before decellularization (Figure 1C/D). Thus, these results indicated that the decellularization process did not cause severe compositional damages to the PDLSC-ECM.

**Figure 1:**
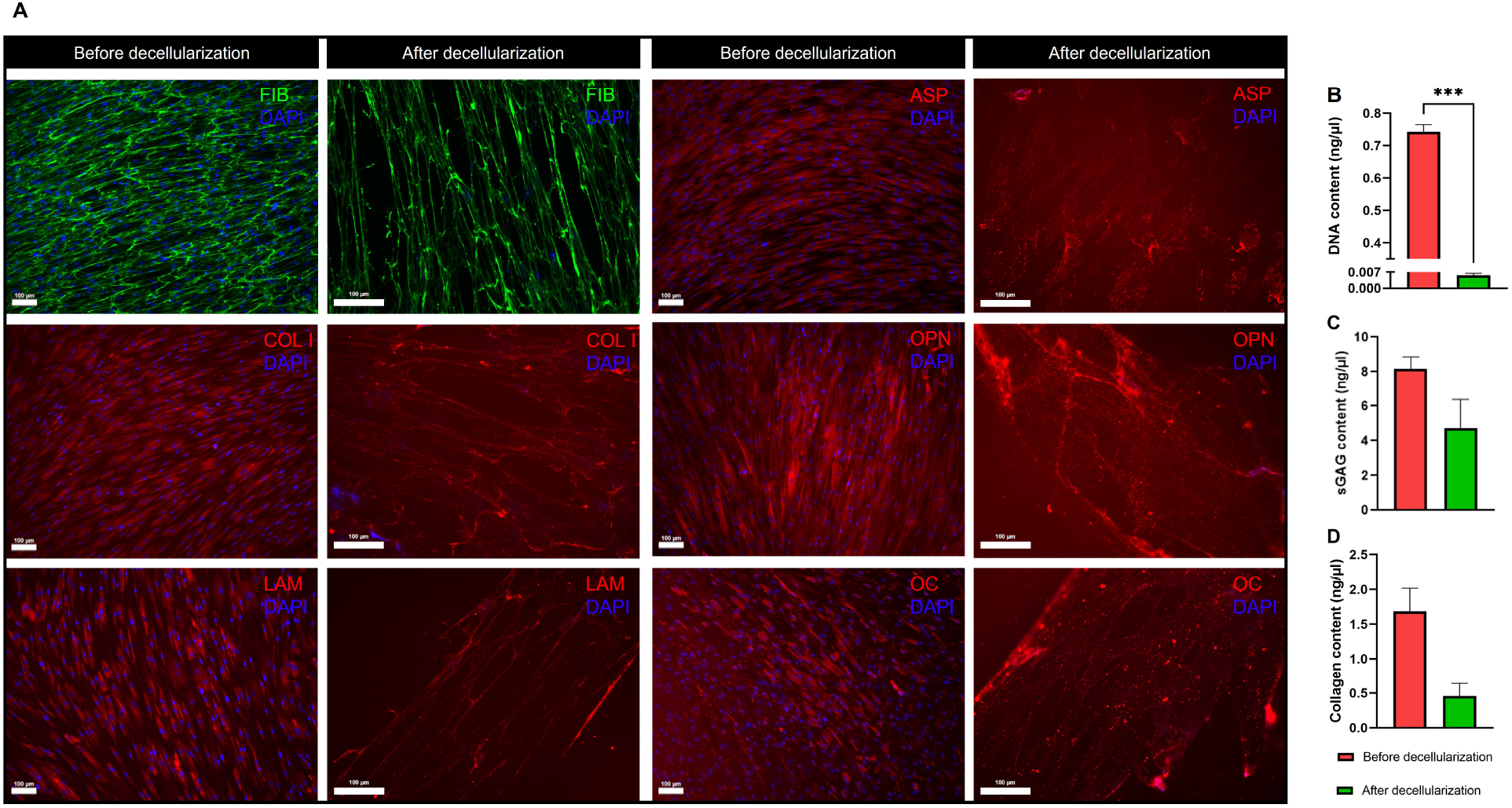
Characterization of decellularized extracellular matrix (dECM) derived from human periodontal ligament stem cells (PDLSCs). A) Immunofluorescence staining images of fibronectin (FIB, green), collagen I (COL I, red), laminin (LAM, red), asporin (ASP, red), osteopontin (OPN, red) and osteocalcin (OC, red) before and after decellularization DAPI staining (blue) revealed the absence of nuclei after decellularization. Scale bar 100 μm. B) DNA content present in PDLSCs and dECM. C) Quantification of sulphated glycosaminoglycans (sGAGs) in PDLSCs and dECM. D) Content of collagen present in PDLSCs and dECM. Three different samples were used in each quantification assay (N=3) for both conditions; *** p < 0.001.

### 3.2. Characterization of Cell-derived ECM Electrospun Scaffolds

#### 3.2.1. Scanning Electron Microscopy and Energy Dispersive X-Ray Analysis

SEM micrographs of PCL, PCL-CTS and PCL-CTS-ECM electrospun fibers showed that all the scaffolds were composed of beadless and homogeneous nanofibers (Figure 2A, B, C). However, PCL electrospun scaffolds were more heterogenous in terms of fiber diameter, but were still constituted of beadless fibers in the nanometer range (Figure 2A). All scaffolds were highly porous and showed high interconnectivity. dECM particles were clearly detected on top of the PCL-CTS-ECM electrospun fibers, as illustrated in Figure 2D. The average fiber diameter of electrospun PCL fibers was 284 ± 150 nm, whilst the average fiber diameters of PCL-CTS and PCL-CTS-ECM fibers were 121 ± 27 nm and 127 ± 25 nm, respectively (Figure 2E, F, G). The presence of CTS in the fibers composition led to a significant decrease in the fiber diameter. PCL-CTS and PCL-CTS-ECM fibers presented similar diameters at the nanoscale, indicating that the incorporation of dECM into the PCL-CTS solution did not affect the average diameter of the fibers and the electrospinning process. The results from EDX analysis are summarized in Supplementary Table A.1 and the obtained EDX spectra are presented in Supplementary Figure A.2. EDX analysis showed that carbon and oxygen were the main constituents of the scaffolds (Supplementary Table A.1). PCL-CTS and PCL-CTS-ECM scaffolds presented similar carbon and oxygen percentages, differing slightly from the PCL scaffolds. Both PCL and CTS were composed of carbon and oxygen, however CTS presented also nitrogen in its composition. Nitrogen was only detected in scaffolds containing CTS, namely PCL-CTS and PCL-CTS-ECM scaffolds.

**Figure 2:**
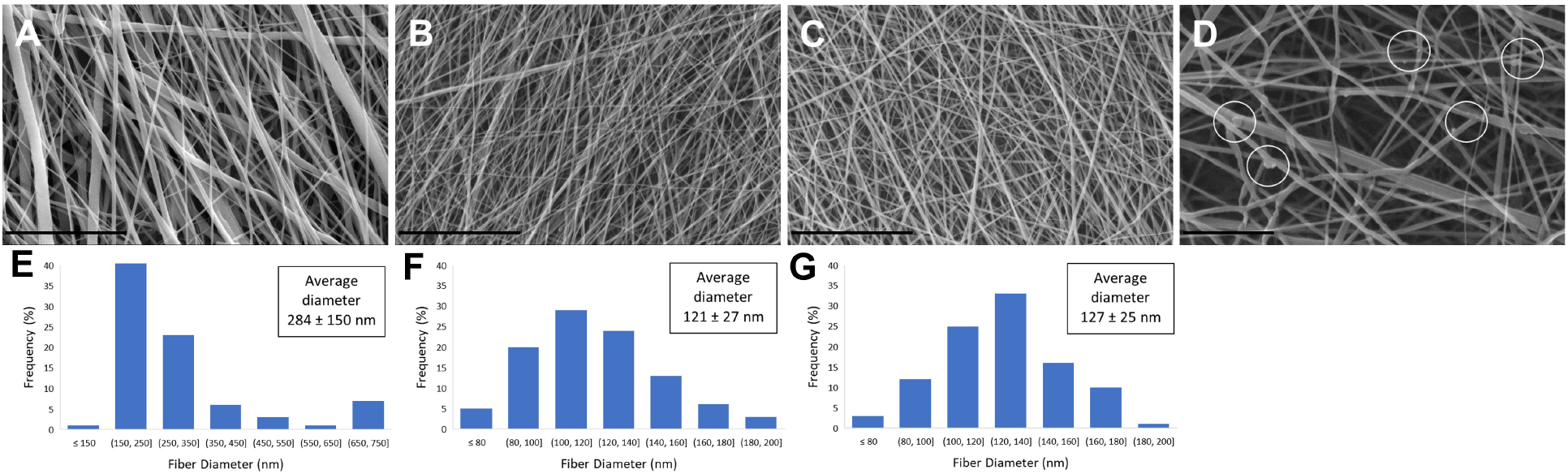
Cell-derived ECM electrospun scaffold characterization. SEM micrographs of PCL (A), PCL-CTS (B) and PCL-CTS-ECM (C) electrospun fibrous scaffolds. Scale bar 8 μm. D) SEM image of dECM particles, identified with white circles, on PCL-CTS-ECM fibers. Scale bar 3 μm. Fiber diameter distribution histograms of PCL (E), PCL-CTS (F) and PCL-CTS-ECM (G) electrospun scaffolds.

#### 3.2.2. Fourier Transform Infrared Spectroscopy

The FTIR spectra of PCL, PCL-CTS and PCL-CTS-ECM electrospun scaffolds showed all the major characteristic IR peaks of PCL (2940, 2865, 1725, 1240 and 1175 cm^-1^). The peaks at 2940 and 2865 cm^-1^ corresponded to asymmetric and symmetric C-H_2_ stretching, respectively. The pronounced peak at 1725 cm^-1^ corresponded to ester carbonyl bond stretching, and peaks at 1240 and 1175 cm^-1^ corresponded to asymmetric and symmetric C–O-C stretching, respectively. The FTIR spectra of CTS and of dECM showed a broad band between 3500 and 3100 cm^-1^ associated with O-H and N-H stretching, and a peak at 1640 cm^-1^ related to C=O stretching of amide I. The broad band resulted in a slight, almost imperceptible deformation in that region in comparison to the PCL scaffold’s spectrum. The peak at 1640 cm^-1^ resulted in a visible deformation around 1670 cm^-1^. Other peaks present in the CTS and dECM spectra are marked as red lines in Figure 3. All the prominent peaks present in the spectra of the used materials (PCL, CTS and dECM) are displayed in Supplementary Table A.2 along with the corresponding vibrations

**Figure 3:**
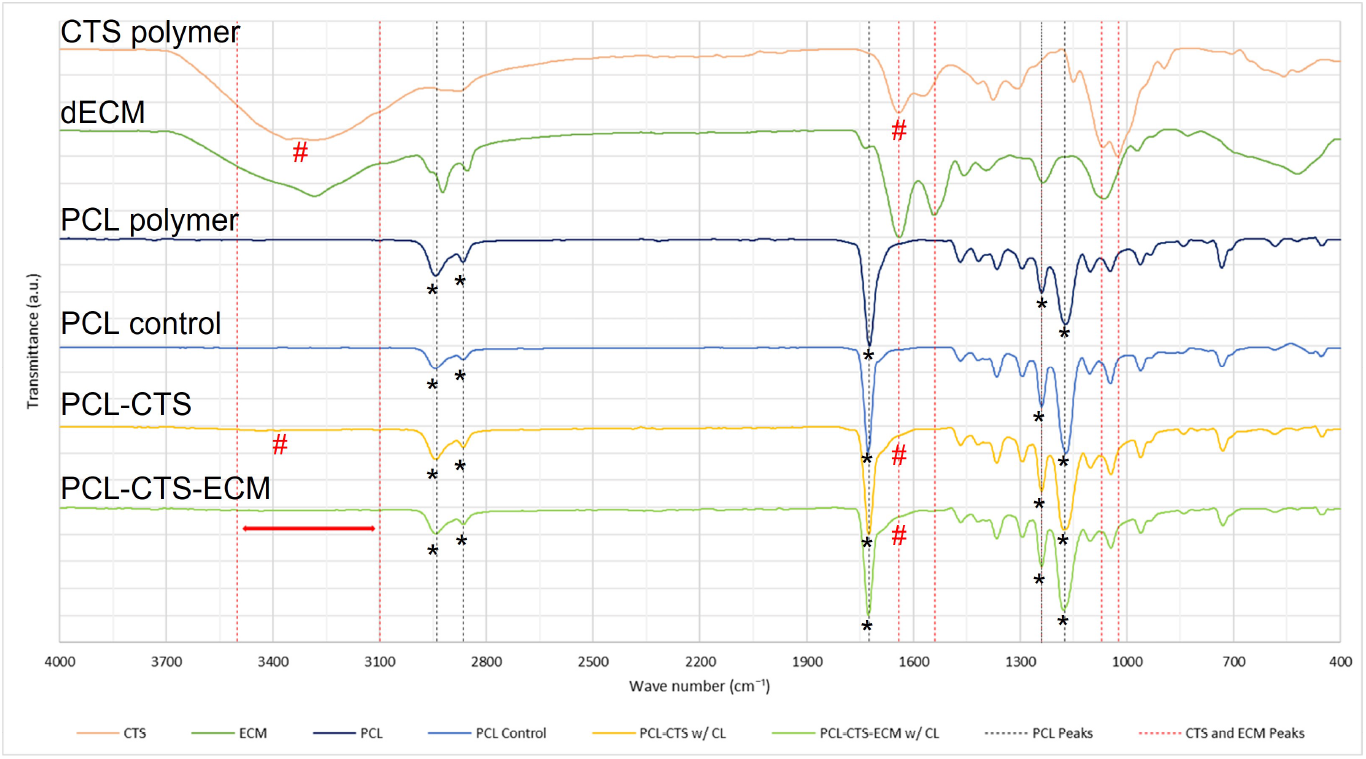
FTIR spectra of lyophilized dECM, PCL and CTS polymers, as well as PCL (PCL Control), PCL-CTS and PCL-CTS-ECM electrospun scaffolds. Asterisks (*) symbols are used to indicate the IR characteristic peaks of PCL. Number signs (#) are used to indicate the IR characteristic peaks of CTS and the resulting deformations.

#### 3.2.3. Thermal Properties Analysis

Thermal properties analysis of cell-derived ECM electrospun scaffolds was performed using differential scanning calorimetry (DSC) and thermogravimetric analysis (TGA). DSC and TGA thermograms of the electrospun scaffolds and the respective pristine polymers, as well as the plots from the first derivative of the mass loss curves (DTGA), are shown in Figure 4 and their thermal properties are summarized in Supplementary Table A.3. PCL, PCL-CTS and PCL-CTS-ECM electrospun scaffolds showed characteristic endothermic (melting) transformation points at 64.84 ± 1.13°C, 64.29 ± 0.29°C and 64.44 ± 0.50°C, respectively. The melting temperatures (T_m_) were similar to the one from the PCL polymer, 65.42 ± 0.50°C (Figure 4A). CTS exhibited a melting temperature of 84.54 ± 1.31°C. The PCL electrospun scaffold had a degradation temperature (T_d_) of 391 °C, which was lower than the value obtained for PCL polymer (410°C). PCL-CTS and PCL-CTS-ECM scaffolds had degradation temperatures of 406°C and 408°C, respectively, which were higher than the PCL scaffold (Figure 4B). CTS showed two degradation steps: a slight degradation at the beginning of the analysis, as can be easily observed in Figure 4B and 4C, and a clear degradation at 298°C. PCL-CTS and PCL-CTS-ECM scaffolds showed an initial degradation step around 250°C, which was not present in the PCL polymer and PCL scaffold. CTS polymer showed a weight loss of 61.53 ± 0.77 %. PCL-CTS and PCL-CTS-ECM scaffolds showed less weight loss than PCL scaffolds, as can be seen in Figure 4B at temperatures above 450°C. The presence of dECM had no significant effect on the thermal properties of the scaffolds.

**Figure 4:**
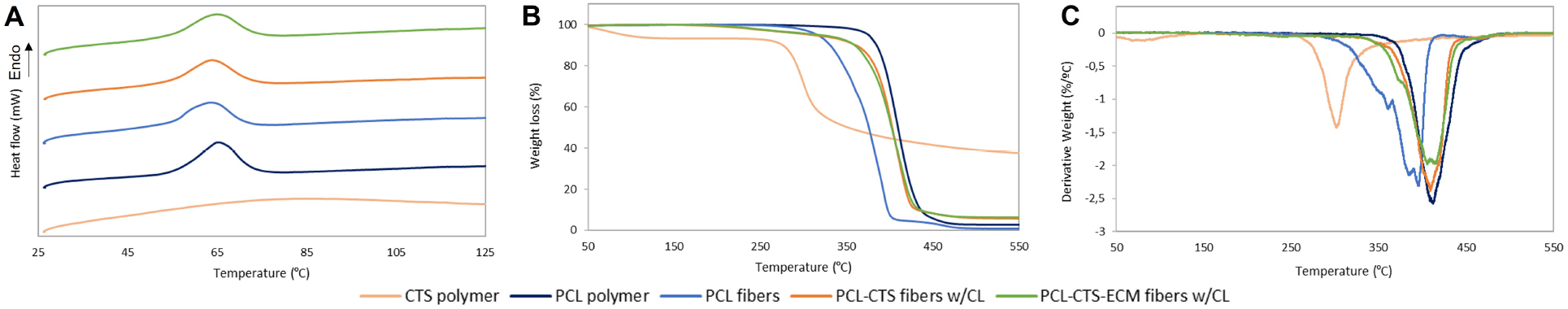
Representative DSC (A), TGA (B) and DTGA (C) curves of PCL and CTS polymers, PCL, PCL-CTS and PCL-CTS-ECM electrospun scaffolds.

#### 3.2.4. Contact Angle

PCL is a hydrophobic polymer, whilst CTS is hydrophilic. PCL scaffolds were hydrophobic, since they presented an average contact angle of 109.2 ± 4.1°. PCL-CTS and PCL-CTS-ECM scaffolds were hydrophilic, since they showed average contact angle values of 46.9 ± 1.6°and 43.8 ± 2.7°, respectively. The addition of CTS to the scaffolds resulted in a significant decrease of the contact angle and increase of hydrophilicity in comparison to the PCL scaffold. Images taken upon droplet placement on the scaffolds illustrate the difference between the hydrophobic PCL scaffold and the hydrophilic CTS-containing scaffolds (Supplementary Figure A.3).

#### 3.2.5. Mechanical Tensile Testing

Results of mechanical tensile testing showed that PCL-CTS and PCL-CTS-ECM scaffolds exhibited decreased elastic modulus, ultimate tensile strength and elongation compared to PCL scaffolds (Figure 5 and Supplementary Table A.4). The addition of CTS in the electrospun scaffolds led to a statistically significant decrease of the mechanical properties due to its brittle behaviour. PCL-CTS and PCL-CTS-ECM exhibited similar mechanical properties. Thus, the incorporation of ECM into the PCL-CTS solution did not drastically affect the mechanical properties of the electrospun scaffolds.

**Figure 5:**
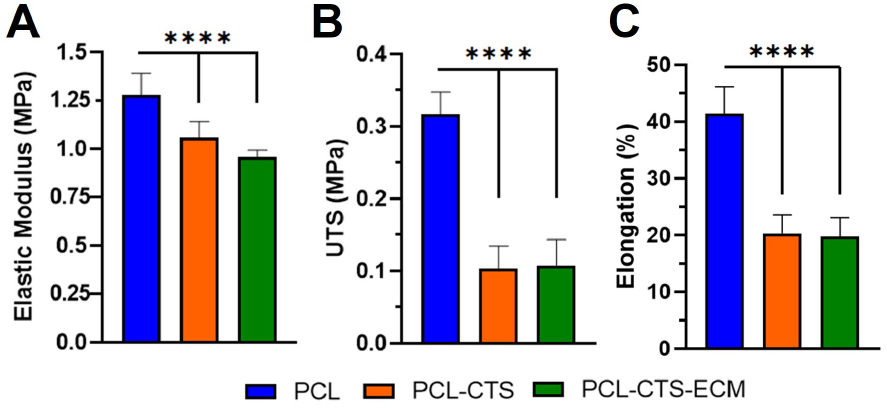
Mechanical properties of PCL, PCL-CTS and PCL-CTS-ECM electrospun scaffolds obtained after uniaxial mechanical tensile testing: elastic modulus (A), ultimate tensile strength (UTS) (B), and ultimate elongation (C). Values are expressed as mean ± SD. Ten different sample specimens (N=10) were used in the analysis; **** p < 0.0001.

### 3.3. Biological Performance of Cell-derived ECM Electrospun Scaffolds

#### 3.3.1. PDLSCs Proliferation Assay and Cell Morphology Assessment

Metabolic activity of PDLSCs was measured on days 1, 7, 14 and 21 to evaluate the effect of cell-derived ECM electrospun scaffolds on PDLSC proliferation (Figure 6A). Interestingly, at days 1 and 7, PCL scaffolds showed a significant increase in cell number compared to PCL-CTS and PCL-CTS-ECM scaffolds. However, after 21 days, PCL-CTS and PCL-CTS-ECM scaffolds presented a statistically significant higher fold increase in the number of cells (Figure 6B), in comparison to PCL scaffolds. At days 14 and 21, PCL-CTS and PCL-CTS-ECM scaffolds showed a statistically significant increase in cell number compared to PCL scaffolds. PCL-CTS-ECM scaffolds showed a statistically significant increase in cell number at days 7, 14 and 21, and significantly higher fold increases compared to PCL-CTS scaffolds. Thus, the cell number increase suggests a beneficial PDLSCs response to the presence of dECM in the composition of the electrospun fibers. After 21 days under osteogenic differentiation conditions, the morphology of PDLSCs cultured on the different electrospun scaffolds was assessed by DAPI-Phalloidin staining (Figure 6C-E). PDLSCs seeded on all electrospun scaffolds presented similar morphology and were densely organized and similarly distributed across all the scaffolds’ structure.

**Figure 6:**
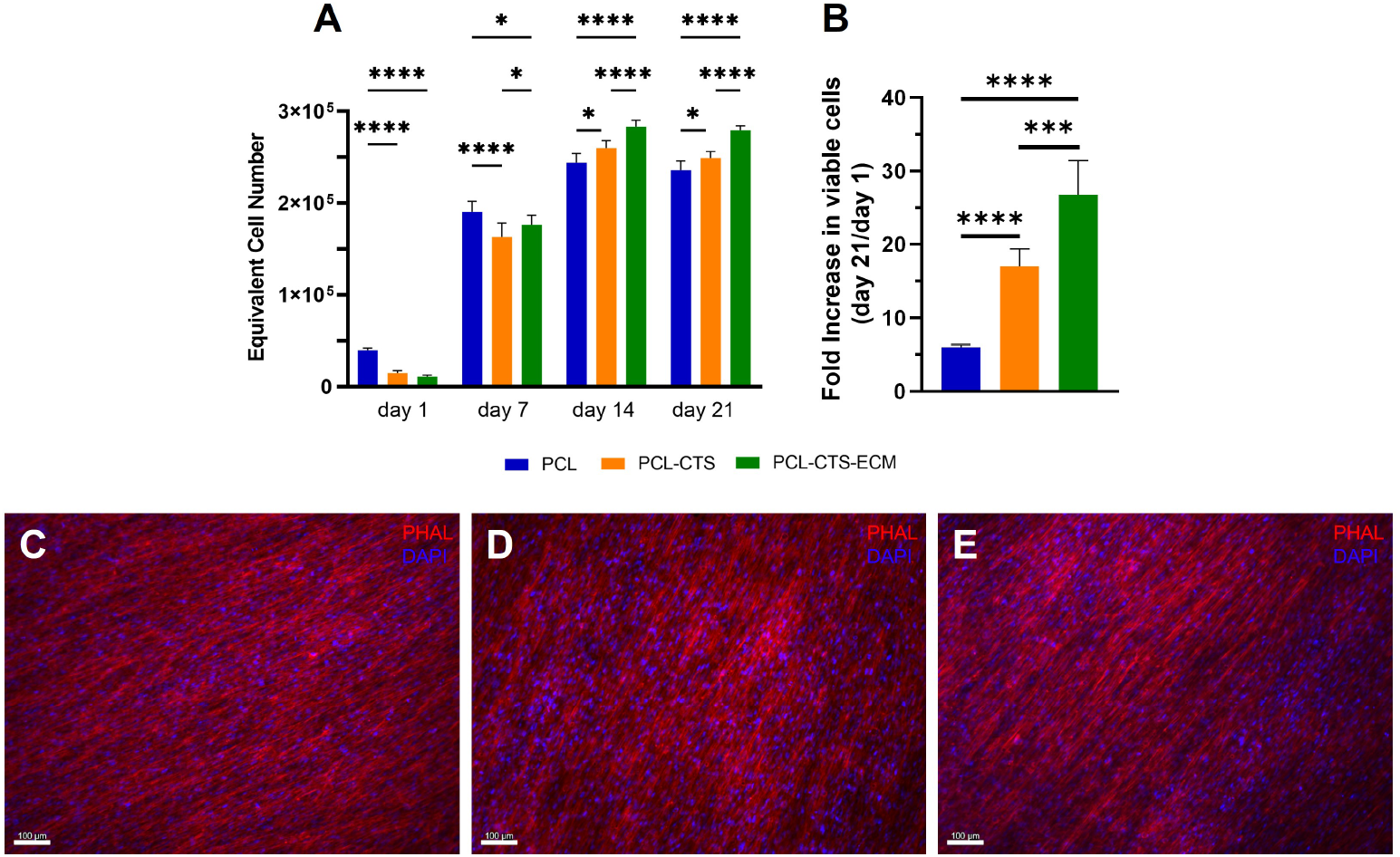
Effects of cell-derived ECM electrospun scaffolds on PDLSCs proliferation. PDLSCs numbers at days 1, 7, 14 and 21 (A) and fold increase (B) in the number of viable cells at day 21 (in relation to day 1) on PCL, PCL-CTS and PCL-CTS-ECM electrospun scaffolds. For each condition, six different samples (N=6) were considered in the analysis; * p < 0.05, *** p < 0.001, **** p < 0.0001. Cell morphology assessment by DAPI-Phalloidin (PHAL) staining at day 21 on PCL (C), PCL-CTS (D) and PCL-CTS-ECM (E) electrospun scaffolds. Scale bar 100 μm.

#### 3.3.2. PDLSCs Mineralization and Osteogenic Differentiation

After 21 days of osteogenic differentiation, SEM micrographs showed that PDLSCs were densely populating the surface of all electrospun scaffolds (Figure 7), consistent with the results from the cell proliferation assay and DAPI-Phalloidin staining (Figure 6) Additionally, elemental analysis of PDLSCs differentiated for 21 days on the different electrospun scaffolds was performed to confirm the presence of calcium and phosphorous, resulting from the cell mineralization process (Figure 7 G-I). The results of the EDX analysis are summarized in Supplementary Table A.5. Calcium and phosphorous were detected on all the scaffolds after 21 days of osteogenic differentiation.

**Figure 7:**
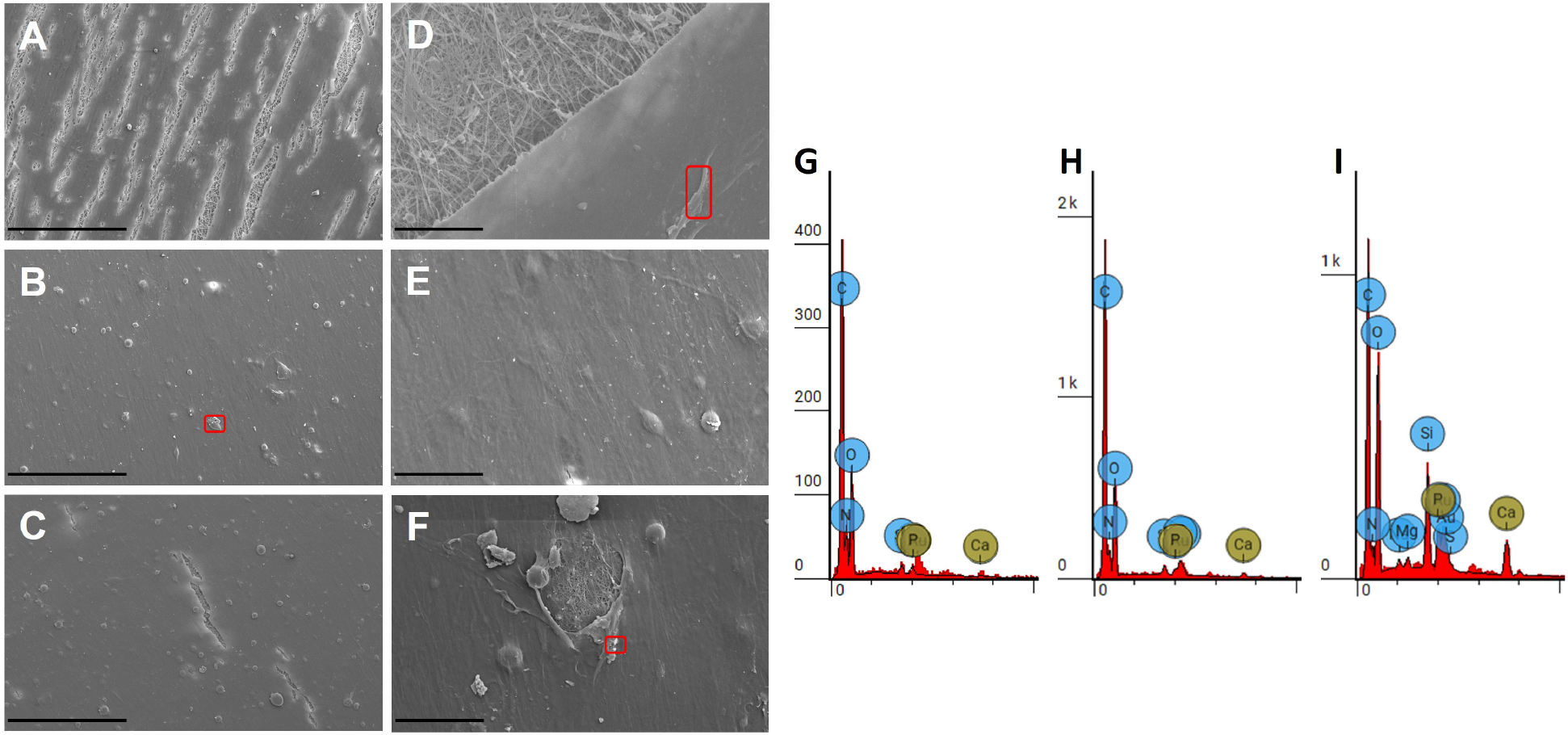
Morphology and elemental composition analysis of PDLSCs cultured on PCL, PCL-CTS and PCL-CTS-ECM electrospun fibers after 21 days under osteogenic differentiation conditions. SEM images of PDLSCs on PCL (A,D), PCL-CTS (B,E) and PCL-CTS-ECM (C,F) electrospun scaffolds, after 21 days of osteogenic differentiation. Spots in which EDX analysis was performed are outlined in red. Scale bars 200 μm (A, B, C) and 30 μm (D, E, F). EDX spectrogram for PCL (G), PCL-CTS (H) and PCL-CTS-ECM (I) electrospun scaffolds after 21 days of osteogenic differentiation.

PDLSCs cultured on PCL-CTS and PCL-CTS-ECM electrospun scaffolds presented a stronger ALP staining compared to the ones cultured on PCL scaffolds, with a more reddish color (Figure 8A). Furthermore, after 21 days of osteogenic differentiation, PDLSCs cultured on PCL-CTS-ECM scaffolds presented significantly higher ALP activity values compared to PDLSCs cultured on PCL and PCL-CTS scaffolds (Figure 8B).

**Figure 8:**
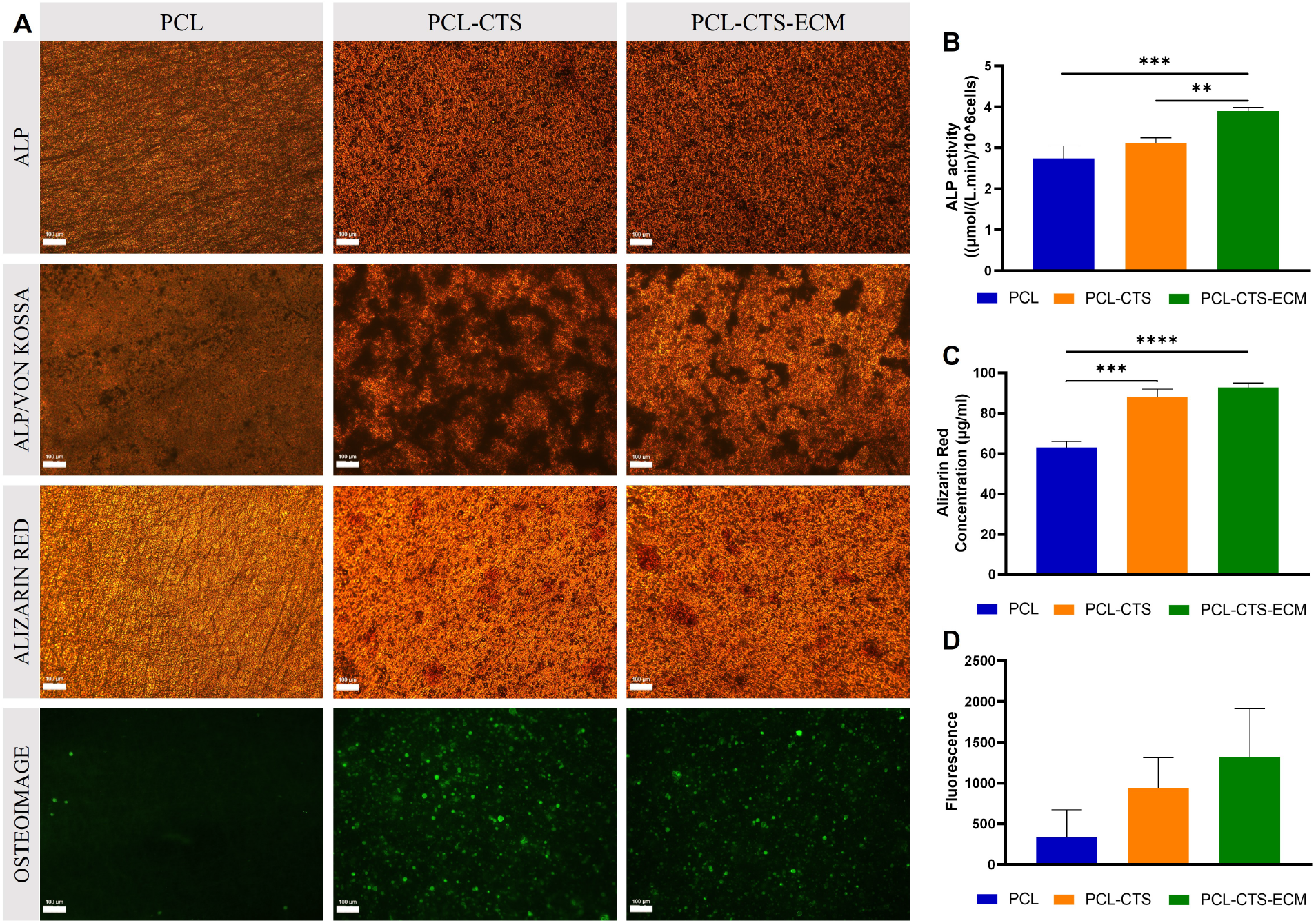
Osteogenic differentiation and mineralization of PDLSCs cultured on PCL, PCL-CTS and PCL-CTS ECM electrospun scaffolds. A) ALP, ALP/Von Kossa, Alizarin Red and OsteoImage stainings of PDLSCs cultured on PCL, PCL-CTS and PCL-CTS-ECM scaffolds after 21 days under osteogenic differentiation conditions. Scale bar 100 μm. B) ALP activity quantification normalized to the number of cells present on PCL, PCL-CTS and PCL-CTS-ECM scaffolds. C) Quantification of Alizarin Red staining on the different electrospun scaffolds. D) Quantification of cell mineralization expressed as fluorescence intensity values on the different scaffold conditions. Values are expressed as mean ± SD. For each experimental group, three different samples (N=3) were used in the analysis; **p < 0.01, *** p < 0.001, **** p < 0.0001.

Regarding mineralization, Von Kossa stainings demonstrated that PDLSCs cultured on PCL-CTS and PCL-CTS-ECM scaffolds showed the presence of more abundant black mineralized deposits than PDLSCs cultured on PCL scaffolds (Figure 8A). Alizarin Red stainings confirmed the presence of calcium deposition (in red) on PCL, PCL-CTS and PCL-CTS-ECM electrospun scaffolds. Interestingly, PDLSCs cultured on PCL-CTS and PCL-CTS-ECM electrospun scaffolds seemed to present higher amount of calcium deposits, which were not present on PCL scaffolds. Moreover, Alizarin Red quantification confirmed a significant increase in calcium deposition from PDLSCs cultured on PCL-CTS and PCL-CTS-ECM scaffolds compared to PCL scaffolds (Figure 8C).

OsteoImage staining images demonstrated cell mineralization on all electrospun scaffolds. However, PDLSCs cultured on PCL-CTS and PCL-CTS-ECM scaffolds presented a more intense green fluorescence in comparison to cells cultured on PCL scaffolds. The quantification of the fluorescent staining confirmed higher levels of cell mineralization on PCL-CTS and PCL-CTS-ECM scaffolds in comparison to PCL scaffolds (Figure 8D). Furthermore, the incorporation of ECM on the PCL-CTS electrospun scaffolds led to an increase in fluorescence regarding cell mineralization, indicating a benefitial effect of dECM in osteogenic differentiation of PDLSCs.

Different osteogenic marker genes were analyzed (RUNX2, OSX, OC, ALP and COL I), as well as CMP1, which is frequently expressed in periodontal tissues (Figure 9). Compared to the control sample (undifferentiated PDLSCs at day 0), all scaffolds showed upregulated expression of RUNX2 and OSX genes, which are key genes involved in osteogenic differentiation. PCL scaffolds showed increased upregulation of RUNX2 and OSX gene expression. All scaffolds showed similar upregulated OC gene expression. ALP gene expression was downregulated in PCL electrospun scaffolds, however PDLSCs cultured on PCL-CTS-ECM scaffolds significantly upregulated ALP gene expression. The expression of COL I gene was significantly upregulated in PCL-CTS and PCL-CTS-ECM scaffolds, whilst in PCL scaffolds it was similar to the control. Interestingly, the incorporation of ECM in PCL-CTS electrospun scaffolds promoted a statistically significant upregulation of ALP and COL I gene expression. Regarding CMP gene expression, all scaffolds demonstrated a statistically significant decrease compared to the control (PDLSCs at day 0).

**Figure 9:**
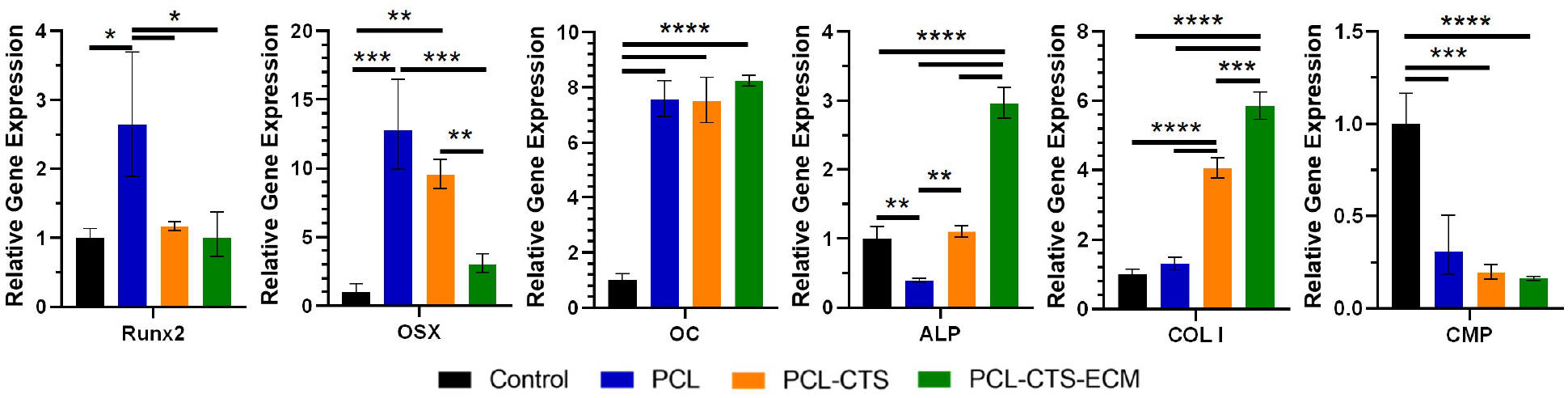
Effects of cell-derived ECM electrospun scaffolds on RUNX2, OSX, OC, ALP, COL I and CMP gene expression by PDLSCs after 21 days of osteogenic differentiation. Gene expressions were normalized to GAPDH gene expression and are presented as fold-change expression relative to the baseline expression of the control sample (undifferentiated PDLSCs at day 0). Values are expressed as mean ± SD (N=3). * p < 0.05, ** p < 0.01, *** p < 0.001, ****p < 0.0001.

The expression of several relevant proteins by PDLSCs on PCL, PCL-CTS and PCL-CTS-ECM electrospun scaffolds after 21 days of osteogenic differentiation is illustrated in Figure 10. Overall, PDLSCs seeded on all types of scaffolds showed a positive expression for collagen I, asporin, osteopontin, osteocalcin, periostin and cementum protein 1.

**Figure 10:**
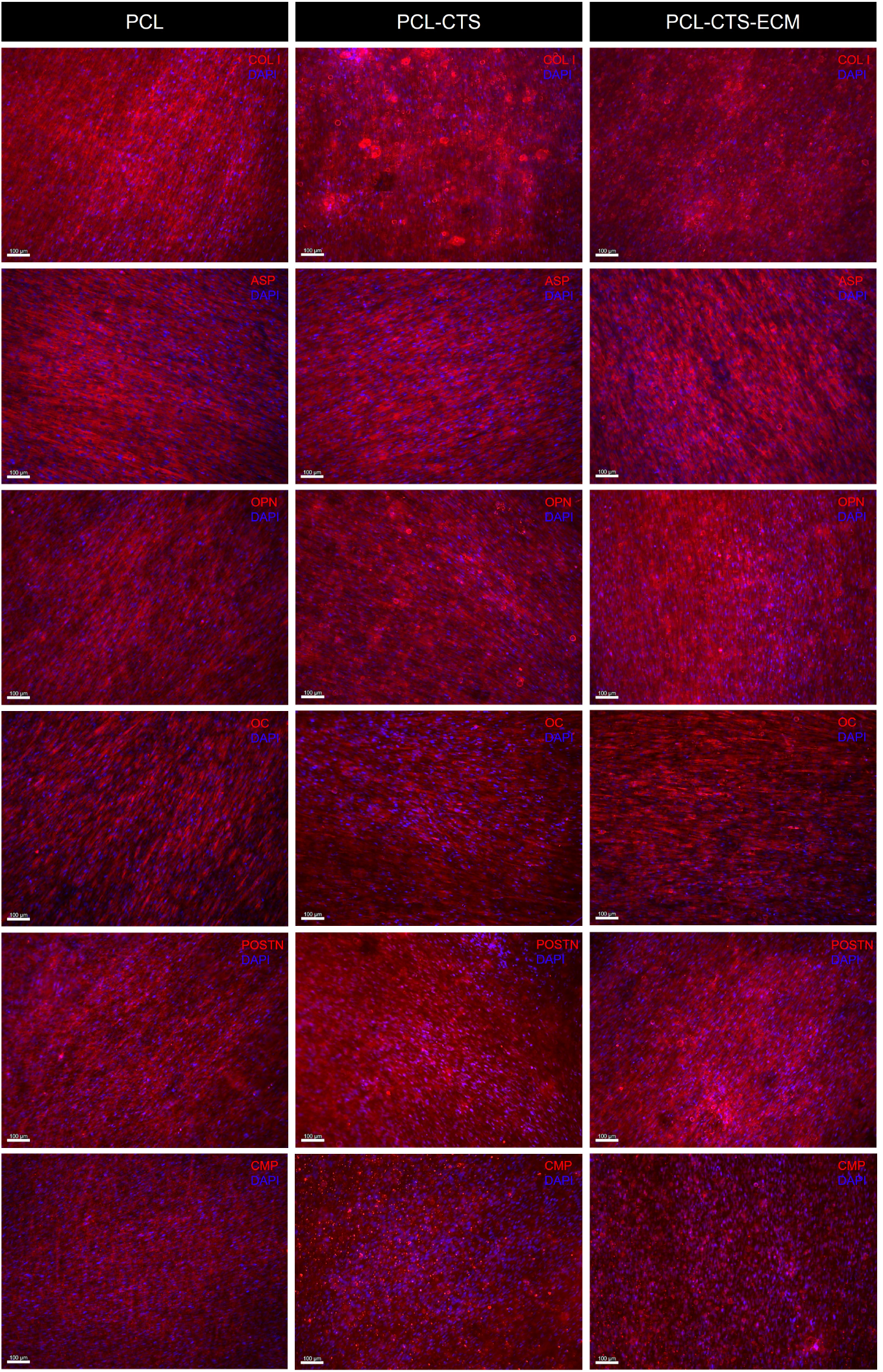
Immunofluorescent staining images of collagen I (COL I, red), asporin (ASP, red), osteopontin (OPN, red), osteocalcin (OC, red), periostin (POSTN, red) and cementum protein 1 (CMP, red) expressed by PDLSCs cultured on PCL, PCL-CTS and PCL-CTS-ECM electrospun scaffolds for 21 days under osteogenic differentiation conditions. Nuclei were counterstained with DAPI (blue). Scale bar 100 μm.

## 4. Discussion

Current periodontal treatments can delay the progression of periodontitis, however if not early treated, periodontitis can progress and will result in destruction of PDL, cementum and alveolar bone [2]. Thus, the ultimate goal of periodontal treatment is to regenerate the whole periodontium. Since the periodontium is a very complex tissue, including soft (PDL) and hard tissues (alveolar bone and cementum), biomaterials used in Periodontal TE should be able to mimic the structure and composition of the tissue and to stimulate the repair of all periodontal tissues. Recently, decellularized cell-derived ECM has been explored as a source of complex molecules that promote cell proliferation and osteogenic activity [20, 22, 23]. However, the ability of cell-derived dECM to induce periodontal regeneration has not been extensively studied. Cell-derived dECM can be used to obtain scaffolds with increased bioactivity and with complexity closer to native tissue microenvironment [25, 27, 29]. In fact, the effect of dECM derived from PDLSCs incorporated in electrospun fibers has not been evaluated for periodontal tissue regeneration. This work describes the first use of lyophilized cell-derived dECM loaded electrospun scaffolds for periodontal TE applications, as a promising therapeutic strategy for periodontitis treatment.

Cell source is known to be an important factor to be considered for the composition of the cell-derived dECM, since cells isolated from different tissues typically produce ECM that mimics the composition of the native tissue matrix [32, 33]. Therefore, taking into account the periodontal microenvironment, in this study we developed cell-derived ECM PCL-CTS electrospun scaffolds composed with dECM derived from PDLSCs.

First, PDLSCs were cultured *in vitro*, until confluence to secrete ECM. Afterwards, a decellularization treatment was applied to remove the cellular components, while retaining the ECM structure. PDLSCs presented a fibroblastlike morphology, whilst PDLSC-derived dECM showed a fibrillary structure, similar to previously reported studies [20, 22, 31]. Regarding immunocytochemistry analysis, PDLSCs expressed collagen I, fibronectin, laminin, asporin, osteopontin and osteocalcin. Interestingly, the decellularization process did not affect the expression of these proteins, which is in accordance with previous works [26, 34]. Type I collagen and fibronectin are two of the predominant proteins present in the native PDL tissue [8]. In fact, previous studies have already reported that collagen I, fibronectin and laminin are main ECM proteins present in cell-derived ECM from various sources [20, 30].

Asporin is an ECM protein, also known as PDL-associated protein 1 (PLAP1) and is predominantly expressed in the PDL [35, 36]. Interestingly, PDLSCs expressed proteins typically present in bone, such as osteopontin and osteocalcin. These two proteins are present in the cementum and in the alveolar bone [8]. Both proteins are osteogenic markers and showed positive expression by PDLSCs, even though these cells were not cultured under osteogenic differentiation conditions. This positive expression demonstrated the intrinsic osteogenic potential of PDLSCs, which *in vivo*, act as a source for renewable progenitor cells that can differentiate into osteoblasts.

After decellularization treatment, no cell nuclei were visible by DAPI staining and DNA content was almost completely diminished, confirming successful decellularization process. Furthermore, sGAGs and collagen contents were retained after decellularization, which demonstrated that the decellularization process did not greatly affect the ECM composition. In fact, besides sGAGs and collagen [26, 31, 37], other growth factors have also been reported to be retained after decellularization treatment [26].

In this work, the electrospun scaffolds fabricated were composed by nanofibers, mimicking the scale of fibers and fibrils present in the native PDL, which are of nano-to microsized order [6, 38]. The presence of CTS in the fibers led to a significant decrease in the fiber diameter, as it has been previously reported in the literature [39–42]. PCL-CTS and PCL-CTS-ECM fibers presented similar diameters, which indicated that the incorporation of dECM did not influence the electrospinning process or altered the average fiber diameter of scaffolds, which was in accordance with previous studies [25]. Nitrogen content detected by EDX analysis confirmed the presence of CTS in PCL-CTS and PCL-CTS-ECM scaffolds. Nitrogen content was slightly higher in PCL-CTS-ECM scaffolds compared to PCL-CTS scaffolds. This difference might be due to the incorporation of dECM, which is composed of proteins, a known source of nitrogen. SEM images of PCL-CTS-ECM electrospun fibers confirmed the presence of cell-derived dECM fragments on the top of the fiber surface, which is concordant with previous works [25, 28].

The FTIR spectra of PCL, PCL-CTS and PCL-CTS-ECM electrospun scaffolds showed all the major characteristic peaks of PCL, which have been extensively reported in the literature [39–41, 43, 44]. The presence of CTS in PCL-CTS and PCL-CTS-ECM scaffolds was confirmed through deformations of certain spectra regions in comparison to the PCL scaffold spectra. Deformations at 3400 cm^-1^ resulting from the broad band in that region and at 1670 cm^-1^ due to CTS presence in the scaffolds have been previously reported in the literature [40–42, 44–46]. Lyophilized dECM powder derived from PDLSCs was also characterized by FTIR. FTIR spectra of lyophilized PDLSC-derived dECM demonstrated some similarities with spectra from other cell-derived ECMs, such as human umbilical vein endothelial cell- and mesenchymal stem/stromal cell-derived ECMs [25, 30]. However, the presence of dECM could not be confirmed by FTIR analysis due to the overlap of the ECM’s specific peaks with those from CTS and PCL. It is also important to note that a low amount of dECM was present in the PCL-CTS-ECM scaffolds composition in comparison to the large amounts of PCL and CTS, therefore no visible differences resulting from the dECM incorporation could be observed, as reported in a previous study from our group [25].

Regarding thermal analysis, the electrospun scaffolds showed characteristic endothermic transformation points at temperatures identical to the PCL polymer [39, 40, 42]. PCL-CTS and PCL-CTS-ECM scaffolds showed an initial degradation step and less weight loss compared to PCL scaffolds, which is in agreement with the existing literature [39, 45, 47] and confirms the presence of CTS. The presence of dECM had no significant effect on the thermal properties of the scaffolds, which is in accordance with previous studies from our group [25].

Contact angle measurements demonstrated that PCL scaffolds were hydrophobic, due to the high contact angle value [40, 42, 43]. The addition of CTS resulted in a decrease of the contact angle, thus enhancing the hydrophilicity of the scaffolds, which might be favorable for cell proliferation. The contact angle values of PCL-CTS and PCL-CTS-ECM were similar to those found in the literature [42, 43]. Interestingly, the incorporation of dECM also did not affect the hydrophilicity of the PCL-CTS electrospun scaffolds, which might be explained by the much lower amount of dECM added in comparison to CTS.

Taking into account that pure CTS fibers have poor mechanical properties and rapid degradation, mechanical tensile testing revealed decreased mechanical properties of PCL-CTS and PCL-CTS-ECM scaffolds in comparison to PCL scaffolds, which is in accordance with previous literature [39, 41, 43, 47]. It has been shown that when the amount of CTS is increased, nanofibers tend to become more brittle [41]. Nevertheless, PCL-CTS and PCL-CTS-ECM scaffolds showed elastic modulus of around 1 MPa, comparable to the modulus of the PDL that ranges between 0.607 and 4.274 MPa under loads between 1 and 5 N [48]. Carvalho and colleagues similarly reported that the incorporation of dECM did not greatly affect the mechanical properties of the electrospun scaffolds [25]. The results demonstrated that PCL-CTS-ECM scaffolds maintained similar physical and mechanical properties of PCL-CTS scaffolds.

The biological performance of the produced PCL-CTS-ECM scaffolds was evaluated in comparison to PCL-CTS and PCL groups by culturing PDLSCs on the different types of scaffolds for 21 days under osteogenic induction conductions. During the 21 days of culture, a statistically significant increase in proliferation of PDLSCs seeded on PCL-CTS and PCL-CTS-ECM scaffolds was observed, in comparison to PCL scaffolds, as confirmed by higher fold increases in the number of viable cells. In fact, PCL-CTS scaffolds have been reported to show increased cell viability and numbers compared to PCL scaffolds [40, 42, 47, 49, 50], due to the high biocompatibility, structural similarity to glycosaminoglycans and hydrophilicity of CTS [14]. Results showed that PCL-CTS-ECM scaffolds presented higher cell numbers compared to the other scaffold groups, which suggests enhanced cell proliferation due to the presence of dECM in the scaffolds as previously reported in the literature [25, 51–53]. Thus, we hypothesized that dECM present in these electrospun fibers may accelerate cell proliferation due to the presence of relevant signalling molecules and growth factors.

Regarding osteogenic differentiation, PDLSCs cultured on PCL-CTS-ECM scaffolds showed significantly higher ALP activity values compared to PCL and PCL-CTS scaffolds. An increase in ALP activity has been reported in the literature in scaffolds with ECM incorporated [25, 51–53]. Furthermore, the presence of CTS in the electrospun fibers also promoted an increase in ALP activity of PDLSCs cultured in PCL-CTS scaffolds in comparison to PCL scaffolds, which has been also previously observed in other studies [49, 50]. In fact, He and co-workers have shown increased ALP activity, extracellular calcium deposition and upregulated OC and RUNX2 gene expression by rat bone marrow mesenchymal cells on PCL-CTS scaffolds compared to pure PCL scaffolds [49]. All the electrospun scaffolds promoted the osteogenic differentiation of PDLSCs, as confirmed by the Alizarin Red and ALP/Von Kossa stainings after 21 days of differentiation. Moreover, Alizarin Red staining and its quantification confirmed significantly increased calcium deposition on PCL-CTS and PCL-CTS-ECM in comparison to PCL scaffolds [49–51]. A more abundant presence of calcium deposits was observed on PCL-CTS and PCL-CTS-ECM in comparison to PCL scaffolds, through the visualization of black precipitates resulting from the Von Kossa staining. The presence of calcium and phosphorous on electrospun scaffolds was further confirmed through elemental analysis. Visualization and quantification of cell mineralization, specifically the hydroxyapatite portion of bone-like nodules deposited by cells, showed higher levels on PCL-CTS and PCL-CTS-ECM in comparison to PCL scaffolds (OsteoImage staining). Notably, these results are in accordance with previous works that have shown that dECM enhances the osteogenic differentiation of MSCs [20, 25, 29]. In particular, Heng and colleagues have shown that PDLSC-derived ECM enhances adhesion, proliferation and osteogenic induction of human dental pulp stem cells, by increasing calcium deposition and ALP activity [22].

Gene expression analysis confirmed the results obtained in the ALP activity assay and ALP staining. ALP gene expression was significantly upregulated in PCL-CTS-ECM group in comparison to the other scaffolds. All scaffolds showed upregulated expression of RUNX2 and OSX gene levels, which are essential genes involved in osteogenic differentiation [54, 55]. Interestingly, the incorporation of dECM in PCL-CTS electrospun scaffolds promoted a statistically significant upregulation of ALP (3-fold increase) and COL I gene expression (5-fold increase). Furthermore, the upregulated expression of OC gene levels, as well as the positive immunofluorescence staining of OPN, OC, POSTN and COL I confirmed the successful osteogenic differentiation of PDLSCs on all electrospun scaffolds.

Regarding periodontal differentiation, PDLSCs cultured on all scaffolds for 21 days showed positive expression of periostin and cementum protein 1. Periostin is a matricellular protein expressed in collagen-rich fibrous connective tissues that are subjected to constant mechanical strains such as the PDL [56]. This protein plays a important role in osteoblast differentiation and survival, therefore positive expression of periostin was expected to be observed in PDLSCs cultured under osteogenic differentiation conditions. Cementum protein 1 is a cementum component, whose presence seems limited to cementoblasts and their progenitors [57]. All scaffolds presented similar downregulated CMP1 gene expression. Reduced expression of CMP1 gene has been reported when PDLSCs were differentiated to osteoblasts *in vitro*, [57]. Undifferentiated PDLSCs (day 0), therefore show an increased gene expression of CMP1 and capacity to differentiate into cementoblasts in comparison to the cells that were cultured under osteogenic differentiation conditions on electrospun scaffolds.

Overall, both CTS and dECM have osteogenic properties and their beneficial effects on PDLSCs osteogenic/ periodontal differentiation were observed by an enhancement of ALP activity, calcium deposition and COL I and ALP gene expression compared to PDLSCs cultured on PCL only scaffolds. In fact, Yang and co-workers have demonstrated that the addition of CTS in PCL fibers increased calcium deposition, ALP activity, and the expression of osteopontin in murine pre-osteoblast cells when compared to pure PCL fibers [50]. Recently, Farag and colleagues have shown that PDLSC-derived ECM enhances osteogenic differentiation, by increasing calcium deposition, ALP activity and expression of osteogenic markers (osteocalcin, osteopontin, bone sialoprotein and bone morphogenetic protein 2), [22, 27]. Regarding periodontal TE applications, decellularized PDLSC-derived ECM can mimic the specific *in vivo*, microenvironment of PDL, including its complex bioactivity, and promotes periodontal regeneration without the potentially immunogenic effects of cellular material. In this work, lyophilized dECM derived from PDLSCs was used to fabricate PCL-CTS-ECM electrospun fibers. Thus, instead of seeding cells on the surface of electrospun scaffolds, allowing the cells to secrete ECM to decorate these scaffolds, the ECM can be produced in culture plates *in vitro*, and further decellularized, collected and lyophilized. This process facilitates the use of dECM which can be easily incorporated in the electrospinning process.

By combining PCL, CTS and dECM, electrospun scaffolds were developed with desirable mechanical properties, enhanced bioactivity and superior osteogenic/periodontal potential. PCL was responsible for providing the mechanical and structural backbone of the electrospun scaffolds. CTS increased the hydrophilicity, enhanced the cell proliferation and promoted osteogenesis, while slightly reduced the mechanical properties. CTS also offers the scaffold a widely reported antibacterial activity, which is particularly relevant in the context of periodontal regeneration settings [58–60]. PDLSC-derived ECM increased the bioactivity and further enhanced proliferation and osteogenic differentiation, without altering the scaffold’s physical and mechanical properties. Overall, PCL-CTS-ECM electrospun fibers promoted PDLSCs proliferation and their osteogenic differentiation *in vitro*, by better mimicking the *in vivo*, ECM composition and structure of PDL. However, further studies such as the analysis of PDLSC-derived ECM composition (e.g., proteomics) and optimization of dECM amounts loaded in the scaffold will be developed to obtain constructs with a optimal performance for periodontal regeneration. Furthermore, *in vivo*, testing of these electrospun scaffolds will also be considered to further assess their potential for periodontal TE applications.

## 5. Conclusions

In summary, we have successfully fabricated and characterized PCL-CTS-ECM electrospun scaffolds with improved bioactivity and native-like mechanical/physical properties for PDL regeneration. Our results showed that cell-derived ECM loaded electrospun nanofibrous PCL-CTS scaffolds enhanced the proliferation and osteogenic differentiation of PDLSCs. PCL-CTS-ECM scaffolds were successfully developed aiming to mimic the structure, architecture, mechanical properties and composition of the native periodontal niche. This work presents, for the first time, the combination of lyophilized dECM derived from PDLSCs with electrospun scaffolds for periodontal TE applications, highlighting its potential for the treatment of periodontitis. Future studies on ECM loaded electrospun scaffolds still need to be conducted to assess their performance *in vivo*, and optimize dECM amounts for improved periodontal tissue differentiation and maturation.

## A. Supplementary Information

**Figure A.1:**
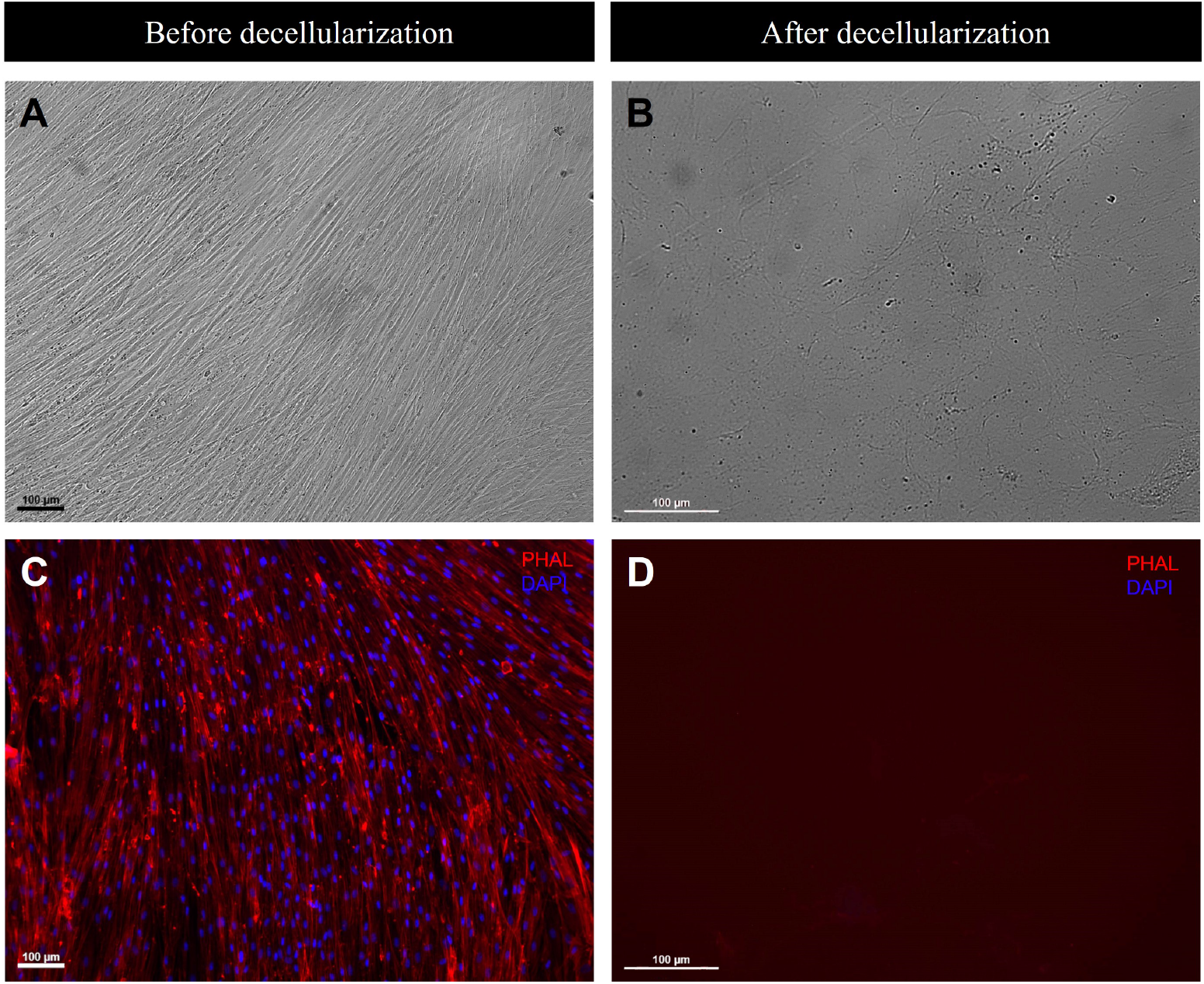
Bright field image of PDLSCs (A) and dECM (B). Cell morphology assessment by DAPI-Phalloidin staining before (C) and after decellularization (D). The cytoskeleton actin filaments were stained with phalloidin (PHAL, red) and nuclei were stained with DAPI (blue). Scale bar 100 μm.

**Figure A.2:**
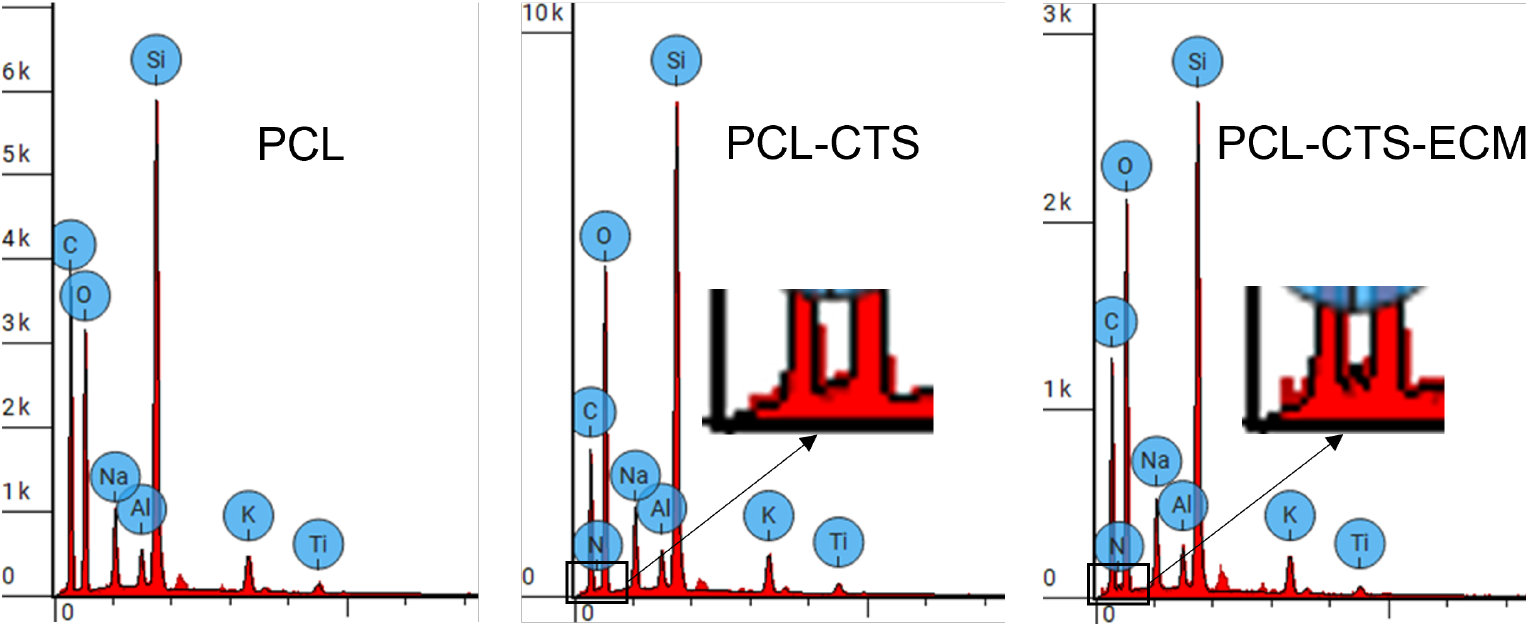
EDX spectra of PCL, PCL-CTS and PCL-CTS-ECM electrospun scaffolds.

**Table A.1:** Atomic percentages of carbon, oxygen and nitrogen of PCL, PCL-CTS and PCL-CTS-ECM scaffolds obtained from EDX analysis.

| Scaffold | C(%) | O(%) | N(%) |
| --- | --- | --- | --- |
| PCL | 57.167 | 27.298 | - |
| PCL-CTS | 39.395 | 38.005 | 1.829 |
| PCL-CTS-ECM | 43.869 | 36.701 | 2.794 |

**Table A.2:**
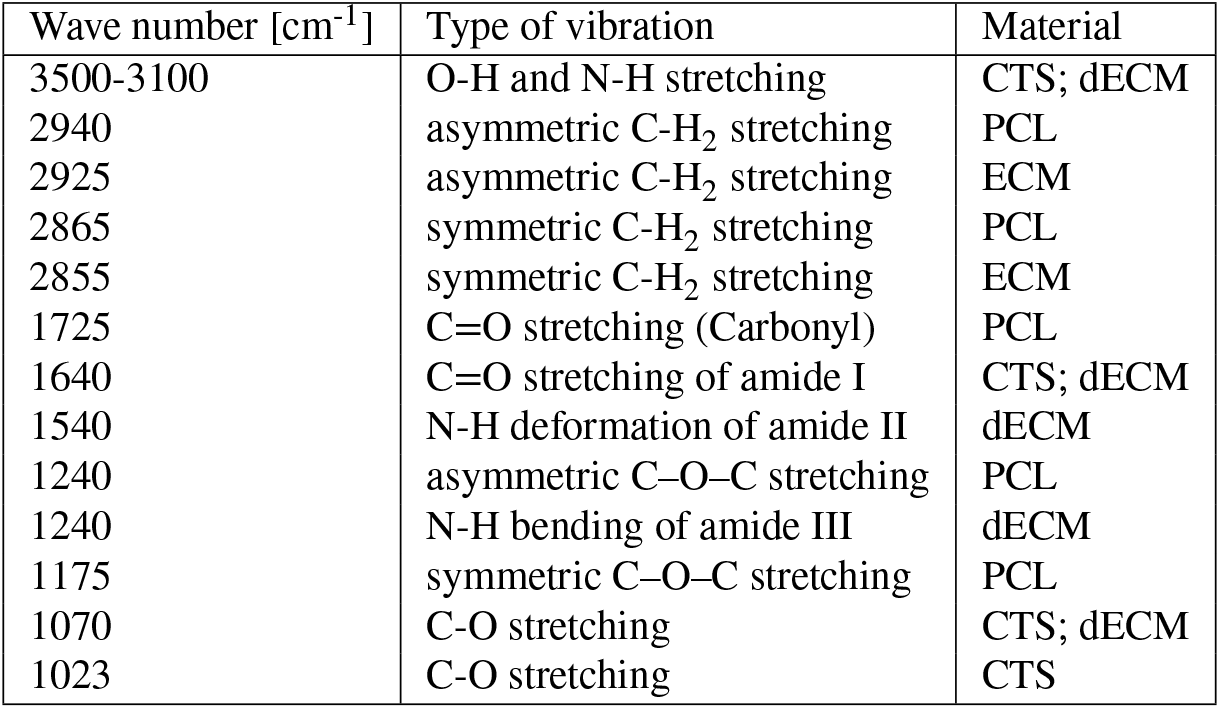
FTIR transmittance peaks and bands present in the spectra of PCL and CTS polymers, and dECM, with corresponding functional groups and types of vibration.

| Wave number [$\text{cm}^{-1}$] | Type of vibration | Material |
| --- | --- | --- |
| 3500-3100 | O-H and N-H stretching | CTS; dECM |
| 2940 | asymmetric C-H<sub>2</sub> stretching | PCL |
| 2925 | asymmetric C-H<sub>2</sub> stretching | ECM |
| 2865 | symmetric C-H<sub>2</sub> stretching | PCL |
| 2855 | symmetric C-H<sub>2</sub> stretching | ECM |
| 1725 | C=O stretching (Carbonyl) | PCL |
| 1640 | C=O stretching of amide I | CTS; dECM |
| 1540 | N-H deformation of amide II | dECM |
| 1240 | asymmetric C-O-C stretching | PCL |
| 1240 | N-H bending of amide III | dECM |
| 1175 | symmetric C-O-C stretching | PCL |
| 1070 | C-O stretching | CTS; dECM |
| 1023 | C-O stretching | CTS |

**Table A.3:** Thermal properties of PCL and CTS polymers, PCL, PCL-CTS and PCL-CTS-ECM scaffolds

|  | PCL polymer | CTS polymer | PCL fibers | PCL-CTS fibers | PCL-CTS-ECM fibers |
| --- | --- | --- | --- | --- | --- |
| T<sub>m</sub> (°C) | 65.42 ± 0.50 | 84.54 ± 1.31 | 64.84 ± 1.13 | 64.29 ± 0.29 | 64.44 ± 0.50 |
| Weight loss (%) | 96.70 ± 0.65 | 61.53 ± 0.77 | 99.83 ± 0.24 | 95.92 ± 2.89 | 93.69 ± 0.45 |
| T<sub>d</sub> (°C) | 410.27 ± 0.33 | 298.30 ± 0.22 | 390.88 ± 5.20 | 406.48 ± 2.52 | 408.10 ± 0.93 |

**Figure A.3:**
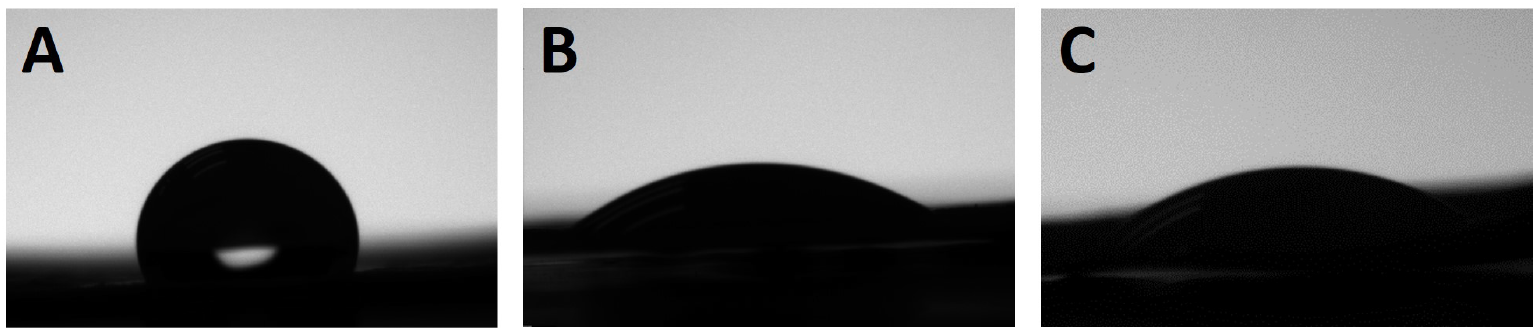
Images from contact angle measurement of PCL (A), PCL-CTS (B) and PCL-CTS-ECM (C) scaffolds.

**Table A.4:** Mechanical properties of PCL, PCL-CTS and PCL-CTS-ECM electrospun scaffolds.

| Scaffold | Modulus (MPa) | UTS (MPa) | Elongation (%) |
| --- | --- | --- | --- |
| PCL | 1.276 ± 0.114 | 0.3172 ± 0.030 | 41.34 ± 4.78 |
| PCL-CTS | 1.057 ± 0.085 | 0.103 ± 0.031 | 20.27 ± 3.24 |
| PCL-CTS-ECM | 0.956 ± 0.038 | 0.108 ± 0.036 | 19.72 ± 3.32 |

**Table A.5:** Atomic (AC) and weight (WC) concentrations of carbon (C), oxygen (O), nitrogen (N), phosphorus (P) and calcium (Ca) obtained through EDX analysis on the spots outlined red in Figure 7 of PCL, PCL-CTS and PCL-CTS-ECM electrospun scaffolds after 21 days of osteogenic differentiation.

| Scaffold | C (%) |  | O (%) |  | N (%) |  | P (%) |  | Ca (%) |  |
| --- | --- | --- | --- | --- | --- | --- | --- | --- | --- | --- |
|  | AC | WC | AC | WC | AC | WC | AC | WC | AC | WC |
| PCL | 46.402 | 40.260 | 23.249 | 26.873 | 28.723 | 29.071 | 0.670 | 1.499 | 0.414 | 1.199 |
| PCL-CTS | 53.288 | 42.700 | 20.890 | 22.300 | 23.215 | 21.700 | 0.484 | 1.000 | 0.860 | 2.300 |
| PCL-CTS-ECM | 45.557 | 30.631 | 25.368 | 22.723 | 16.336 | 12.813 | 2.020 | 3.504 | 4.550 | 10.210 |

